# Geometry of Braided DNA Dictates Supercoiling Partitioning

**DOI:** 10.1101/2024.10.08.617221

**Authors:** Yifeng Hong, Gundeep Singh, Hanjie Wang, Seong ha Park, Michelle D. Wang

## Abstract

DNA’s helical structure requires the replisome to rotate relative to the parental DNA during replication, generating supercoiling that partitions ahead and behind the fork. The influence of DNA substrate geometry on supercoiling partitioning and torsional resistance remains unclear. Here, we have engineered DNA-braiding substrates with end separations found during replication, measured braiding torques using an angular optical trap, and interpreted the results using Monte Carlo simulations. A substrate with small separations readily initiates braiding, suggesting that the fork would rotate during replication, partitioning supercoiling behind the replisome and reducing torsional resistance. In contrast, a substrate with a substantial separation at one end imposes a pronounced ‘torsional barrier’ that hinders braiding initiation, suggesting a lack of fork rotation during replication, so supercoiling would partition ahead of the replisome. Our findings reveal a physical mechanism in which DNA fork geometry could modulate replisome dynamics, manage topological stress, and regulate replication progression *in vivo*.

## Introduction

During DNA replication, a replisome duplicates the parental DNA into two daughter strands. Because of DNA’s helical structure, a progressing replisome must track the DNA helical groove via rotation relative to the DNA substrate ^1–6^. This rotation can potentially intertwine the two daughter strands and thereby prevent chromosome segregation during mitosis ^3,7–13^. In addition, DNA replication inherently creates torsional stress that, in turn, hinders the progression of the replisome ^1,3–5,14^. *In vivo*, these difficulties are partially relieved by the action of topoisomerases. Although topoisomerases are essential in relaxing torsional stress ^2,15–17^, they cannot always keep up with replication ^12,18–25^. Therefore, daughter-strand intertwining and torsional stress accumulation can occur during replication elongation, especially as the replisome approaches termination ^8–11,26,27^. Despite this problem being inherent to replication, we have a limited understanding of the mechanics governing supercoiling partitioning to the daughter strands and daughter-strand intertwining (also referred to as braiding), as well as the extent of torsional stress the replisome experiences.

Previously, we hypothesized that this problem is inherently dictated by the torsional mechanical properties of the single- and double-DNA substrates ^4^. Because torque must be balanced at the replisome, supercoiling partitioning between the front and back of the replisome is solely determined by the torsional resistance of twisting the single-DNA substrate in front of the replisome and the torsional resistance of braiding the double-DNA substrate behind the replisome. The torsional resistance to braiding the double-DNA substrate is expected to critically depend on the separation between the two DNA molecules at each end ^28,29^. *In vivo*, the end separation of the two daughter strands at the replisome might be relatively small, on the order of 20-50 nm, as estimated by the size of the replisome ^30–34,4^. At the end distal to the replisome where the two daughter DNA molecules extend away from the fork, the separation may increase significantly until structural maintenance of chromosomes (SMC) proteins bring the two strands together to a distance of about 35 nm ^35–37^. Thus, elucidating how daughter-strand geometry affects DNA supercoiling partitioning requires direct torque measurements of braiding DNA substrates with defined end separations (20-50 nm). This measurement feat has not been previously attainable.

Previous single-molecule studies selected braids among DNA tethers anchored to surfaces at random locations ^4,28,38–51^ and estimated the end separation by assuming the separations at both ends were identical ^4,28,38–41,44,51^. However, it is very challenging to identify a braid with an end separation < 100 nm, and the equal-separation assumption likely does not hold for most surface-tethered DNA braids. Experiments using multiple optical traps can control the geometry of the two DNA molecules while braiding ^52,53^, but the micron-sized dimensions of the trapping particles limit the DNA end separations to large values. Furthermore, although knowledge of the torque to braid DNA is essential, direct torque measurements are lacking in most previous experimental studies.

In this work, we engineered a DNA braiding substrate with an end separation of about 42 nm at both ends to better mimic the *in vivo* geometry, along with two other braiding substrates with larger end separations for comparison. Using the angular optical trap (AOT), we directly measured the torque required to braid these substrates. Our measurements provided the torque required to initiate the braiding, the torsional stiffness of braiding DNA, and the torque needed to buckle a DNA braid into a plectoneme. To better understand how braiding geometry governs the torsional mechanics of braiding, we performed Monte Carlo (MC) simulations to facilitate the interpretation of our experimental results. The framework laid out in this work should provide insights into how braiding geometry impacts replication-generated torsional stress and the extent of the daughter-strand intertwining during replication.

## Results

### Engineered DNA Braiding Substrates with Defined Geometries

To study DNA braiding under well-defined geometries, we developed methods to engineer two DNA braiding substrates (Methods), namely the ‘O substrate’ (Fig. 1a) and the ‘V substrate’ (Fig. 1b). Each substrate was constructed from a customized plasmid (Methods) and contains two DNA segments of essentially identical lengths for braiding.

**Fig. 1.**
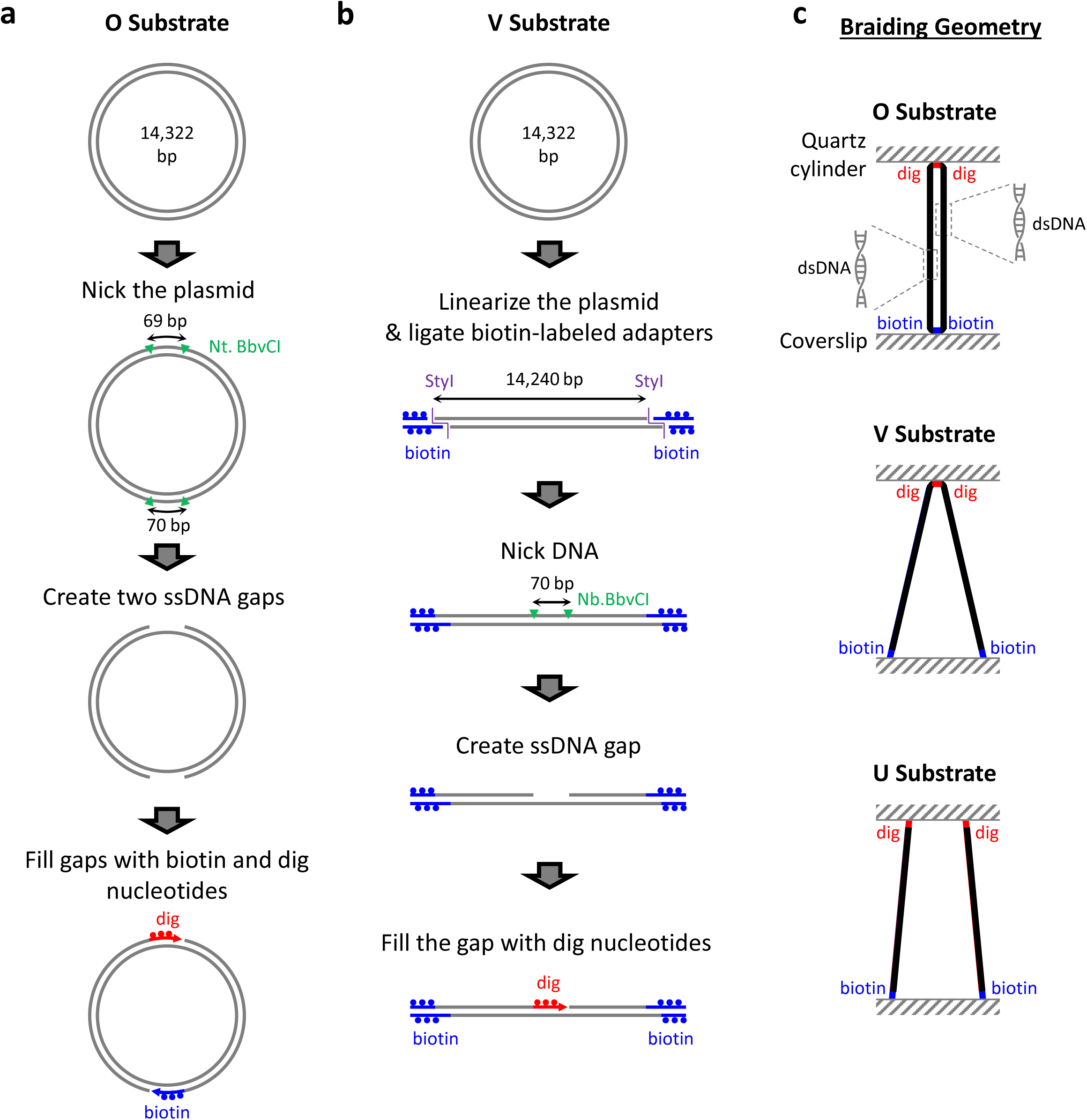
Construction of braiding substrates with defined geometries. **a.** ‘O substrate’ design and preparation. The final product has two dsDNA strands for braiding, each 7.1 kb in length, with minimal separations at both ends. The top end of the ‘O substrate’ has a 69 bp multi-digoxigenin-tagged region (red), while the bottom end has a 70 bp multi-biotin-tagged region (blue). This substrate allows small end separations of the two ‘daughter’ DNA strands once the substrate is anchored between a coverslip and a quartz cylinder held in an AOT. **b.** ‘V substrate’ design and preparation. The final product has two ‘daughter’ strands for braiding, each 7.1 kb in length, with a minimal separation at one end. The center of the substrate contains a 70 bp multi-digoxigenin-tagged region (red). The two biotin-labeled (blue) ends allow variable end separations once anchored to a coverslip surface. **c.** Three different braiding geometries, achieved by different braiding substrates. Each substrate is anchored between the coverslip surface and the bottom surface of a quartz cylinder held by the AOT.

The ‘O substrate’ was designed with minimal separations at both ends (Fig. 1a; Methods). This design best mimics an *in vivo* configuration in which the two replicated DNA strands are held closely together at one end by the replisome and at the other end by SMC proteins. To construct this substrate, we created two ssDNA gaps (69 bp or 70 bp) symmetrically located at half a plasmid length apart, followed by filling in with multi-digoxigenin-labeled or multi-biotin-labeled nucleotides at the two sites. The final product thus has two distinct short surface-anchoring sites that define the end separations. These sites are also multi-tagged, enabling torsional constraint to either an anti-digoxigenin-coated or streptavidin-coated surface. We estimate that each multi-tagged anchoring region yields an effective DNA end separation of approximately 41-47 nm (Methods). This value accounts for the contributions of DNA bending on both sides of the anchoring region and is comparable to that estimated at the replisome. In addition, each ‘daughter’ DNA strand was nicked (Methods) to mimic the *in vivo* configuration where both the leading and lagging strands are thought to be able to rotate around the DNA helical axis during replication ^1,2,4^.

The ‘V substrate’ was designed to have a minimal separation at one end and a greater separation at the other (Fig. 1b; Methods). This design mimics an *in vivo* configuration where daughter strands are held closely together at one end by the replisome, but the other end might be further apart (e.g., when SMC proteins fail to bring the two strands together ^54^). To construct this substrate, the plasmid was linearized and then ligated with a biotin-labeled adapter at each end for surface anchoring. An ssDNA gap was created in the middle of the template, resulting in a 70 bp region containing multi-digoxigenin-labeled nucleotides. Before measurements, each daughter strand in the ‘V substrate’ was nicked to prevent torsion accumulation within each ‘daughter’ DNA (Methods).

Upon surface immobilization between an anti-digoxigenin-coated quartz cylinder optically trapped by the AOT and a streptavidin-coated coverslip surface (Methods), the ‘O substrate’ has a minimal end separation at each end of the two ‘daughter’ DNA molecules (Fig. 1c, top). In contrast, the ‘V substrate’ has a minimal end separation at the quartz cylinder and a variable end separation at the coverslip surface. This variable separation can be substantial compared with the length of each ‘daughter’ DNA (Fig. 1c, middle).

For completeness, we also examined a braiding geometry with wide separations at both ends. We anchored individual 6.5 kb DNA molecules (Methods) at random locations on the coverslip surface and then selected a cylinder that was tethered via two DNA molecules, similar to what has been done previously ^4^. This substrate has substantial separations at both ends compared to the length of each ‘daughter’ DNA molecule (Fig. 1c, bottom). For convenience, we refer to this configuration as the ‘U substrate’. This configuration may not readily occur *in vivo* but we investigated this substrate to allow for direct comparison with prior studies.

### Braiding DNA with Minimal End Separations

The ‘O substrate’ has small end separations at both ends and should best resemble the *in vivo* geometry of daughter strands during replication. It has nearly identical end separations at both ends (*a* and *b*), which are significantly smaller than the length *l* of each ‘daughter’ DNA molecule (*a* = *b* ≪ *l*) (Fig. 2a). To determine the torque required to braid the ‘O substrate’, we used the AOT, which allows simultaneous measurements of torque, rotational angle, force, and extension of DNA via a nanofabricated cylinder ^4,5,55–61^. Once torsionally constrained between the coverslip surface and the cylinder (Fig. 2a; Methods), the ‘O substrate’ was held at a constant force and braided via rotation of the cylinder. The number of turns added by the cylinder equals the catenation number (the number of times the two daughter strands intertwine).

**Fig. 2.**
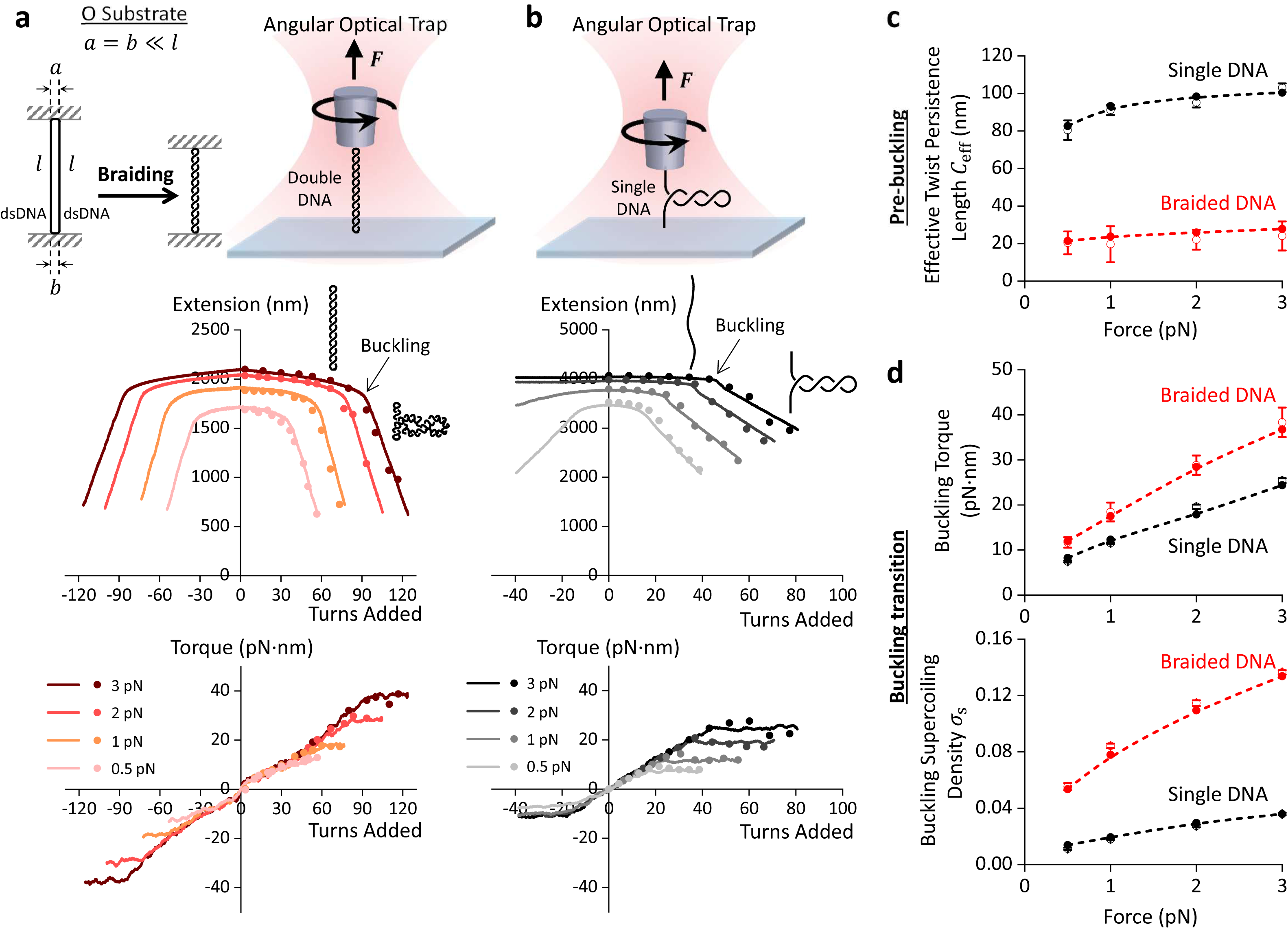
Torsional mechanical properties of braiding the ‘O substrate’, in comparison with those of twisting a single-DNA substrate. **a.** Torsional measurements of braiding the ‘O substrate’ using the AOT. The cartoons at the top show the experimental configuration. The ‘O substrate’ is torsionally constrained between the bottom surface of an optically trapped quartz cylinder and the surface of the coverslip of a sample chamber, creating a configuration with the end separations (*a* and *b*) significantly smaller than the ‘daughter’ DNA length *l*. Both anchoring segments are torsionally constrained to their anchoring surfaces. While turns are added to the substrate via cylinder rotation as the substrate is held under a constant force, extension and torque are simultaneously measured via the AOT. The bottom plots show the measured extension and torque (lines, averaged over *N* = 15 molecules for each force), along with comparison with the MC simulation results (dots). For each force, the measurements are the means of *N* = 15 molecules. Source data are provided as a Source Data file. **b.** Torsional measurements of twisting a single-DNA substrate. The cartoons at the top show the experimental configuration for twisting a single-DNA substrate using the AOT, in a way akin to that of the ‘O substrate’. The single-DNA substrate is 12.7 kb in length, flanked by two end-tagged DNA segments of about 500 bp to torsionally constrain the substrate between the two surfaces (Methods). The bottom plots show the measured extension and torque (lines), along with comparison with the MC simulation results (dots). For each force, the measurements are averaged from *N* = 20, 19, 18, 15 individual molecules for forces at 0.5pN, 1pN, 2pN, and 3pN respectively. Source data are provided as a Source Data file. **c.** Effective twist persistence length *C*_eff_ of braiding the ‘O substrate’ versus force. The *C*_eff_ is obtained from the torque-turns relation near zero turns (Methods) from both measurements (open dots with error bars being SDs) and simulations (solid dots). For comparison, also shown are the corresponding *C*_eff_ data of a single-DNA molecule. The dashed lines are interpolations from the simulation results to guide the eye. For the ‘O substrate’, *N* = 15 individual molecules for each force. For the single substrate, *N* = 20, 19, 18, 15 individual molecules for forces at 0.5pN, 1pN, 2pN, and 3pN respectively. Source data are provided as a Source Data file. **d.** Buckling torque and buckling superhelical density of braiding the ‘O substrate’ versus force. Superhelical density of braided DNA (the catenation density) is defined as *Ca*/*Lk*_0_, where *Ca* is the catenation number and *Lk*_0_ is the linking number of a relaxed DNA for one of the ‘daughter’ DNAs for braiding. Buckling torque and buckling superhelical density are determined from both measurements (open dots with error bars being SDs) and simulations (solid dots). For comparison, also shown are the buckling torque and buckling superhelical density of a single-DNA molecule. The dashed lines are interpolations from the simulation results to guide the eye. For the ‘O substrate’, *N* = 15 individual molecules for each force. For the single substrate, *N* = 20, 19, 18, 15 individual molecules for forces at 0.5pN, 1pN, 2pN, and 3pN respectively. Source data are provided as a Source Data file.

Using the AOT, we measured the torque required to braid the ‘O substrate’ while simultaneously monitoring its corresponding extension (Fig. 2a; Supplementary Fig. 1a). As turns are added, the torque increases due to resistance to braiding, and the extension decreases due to the two ‘daughter’ DNA molecules helically wrapping around each other to form a braided structure. When the braiding torque reaches a critical value, the torque plateaus while the extension shortens sharply, indicating buckling of the braided DNA to form a plectoneme. We found that the number of turns required to buckle a DNA braid increases with force. As more turns are added, the plectoneme of the braided DNA continues to extrude, indicated by a linear extension decrease. These behaviors are reminiscent of the torsional properties of the single-DNA substrate measured under the same conditions (Fig. 2b), consistent with the results from our previous studies ^4,55–64^.

We found that the extension-turns relation is essentially an even function, nearly symmetric about zero turn number, and the torque-turns relation is essentially an odd function, nearly antisymmetric about zero turn number. We detected some slight asymmetries (Supplementary Fig. 2), likely due to helix-specific interactions ^65–69^. These predominately symmetric features suggest that the inherent chirality of DNA molecules does not play a significant role in the torsional mechanical properties of braiding. As our work aims to understand the supercoiling generation during replication, we focus on the torsional mechanics of the right-handed braid (R-braid), which corresponds to (+) supercoiling.

To more rigorously understand these braiding features, we performed MC simulations of the ‘O substrate’ (Fig. 2a) and the single-DNA substrate (Fig. 2b) (Methods; Supplementary Table 1). Although MC simulations have previously been used to study DNA braiding ^28,70^, it was necessary to refine the approach to improve accuracy and achieve quantitative agreement between simulation results and measurements. We therefore extended the model to incorporate a stronger topological invariant to constrain topology and a Debye-Hückel electrostatic interaction potential instead of a hard-wall repulsion between DNA strands (a parameter that plays a key role in determining the braid diameter) (Methods). These extensions are essential at higher catenation numbers, especially after buckling occurs. These improvements enabled us to compute torsional mechanics over a broader range of catenation densities for braided conformations, including torque in the post-buckling regime, which has not been reported in previous MC simulations. We found that the simulated extension-turns and torque-turns relations under different forces agree well with measurements for the ‘O substrate’, providing validation of our simulation method. Notably, the previous MC simulation work on DNA braiding ^28^ predicted a substantially steeper torque-turns relation under the same force than our measurements.

We further analyzed the braiding data shown in Fig. 2a and found that both the measured and simulated torques increase linearly with turns near zero turns, albeit with a slight torque discontinuity due to small but non-zero end separations (Fig. 2a). As more turns are introduced, these torques exhibit some nonlinearity, indicating a twist-stiffening effect (Supplementary Fig. 2a), as suggested by a previous theoretical study ^29^. To characterize the torsional resistance to braiding at a small number of turns, we determined the torsional stiffness of braiding using the slope of the torque-turns relation near zero turns (Methods). We then converted this stiffness to the effective twist persistence length *C*_eff_, a parameter independent of the DNA length ^4^, and found it to be insensitive to tension over the range from 0.5 to 3 pN (Fig. 2c).

In comparison with the single-DNA substrate, the ‘O substrate’ has a *C*_eff_ about 4 to 5 times smaller (Fig. 2c), indicating that ‘O substrate’ is easier to twist. However, the ‘O substrate’ is more resistant to buckling, with buckling occurring at a higher torque and a larger superhelical density (Fig. 2d). Unlike the single-DNA substrate, the ‘O substrate’ extension slope also shows a non-monotonic force-dependent behavior after buckling (Supplementary Fig. 3). These results from the ‘O substrate’ establish the torsional mechanical properties necessary to model supercoiling partitioning during DNA replication.

### Braiding DNA with a Substantial End Separation

The ‘V substrate’ has one end with a small separation (*a* ≪ *l*) and another with a substantially larger separation (*b*), mimicking a case where the replisome brings one end together, but SMC proteins fail to bring the other end together. The ‘U substrate’ has both ends separated by a substantial separation, a configuration that may not readily occur *in vivo*. We have included studies of the ‘U substrate’ for completeness and comparison with previous studies.

Using the AOT, we measured torque-turns and extension-turns relations of the ‘V substrate’ (Fig. 3a; Supplementary Fig. 1b) and the ‘U substrate’ (Fig. 3b; Supplementary Fig. 1c). The measured relations have some similarities with those measured with the ‘O substrate’ (Fig. 2a; Supplementary Fig. 1a) but with significant differences. The most distinguishing features occur near zero turns (more precisely, between -0.5 turns and +0.5 turns). For both substrates, the torque discontinuity near zero turns, which we refer to as the ‘torque gap’ *τ*_gap_ (Methods), becomes prominent (Fig. 3a,b). Beyond the zero turns before buckling, the extension decreases more steeply with turns, which we characterize by the ‘hat-top slope’, and the buckling occurs at a lower number of turns.

**Fig. 3.**
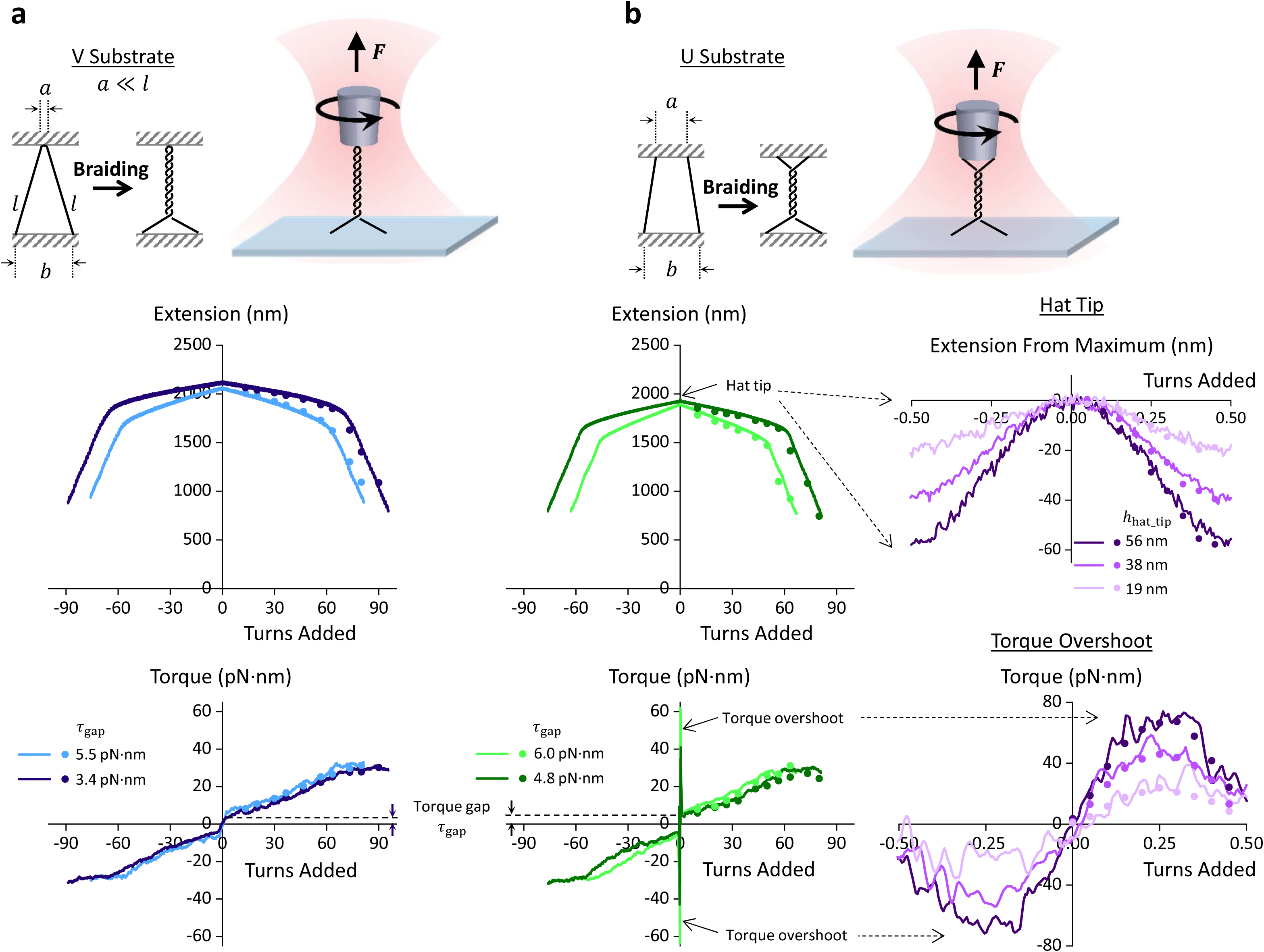
Torsional measurements of braiding the ‘V substrate’ and the ‘U substrate’. **a.** Torsional measurements of braiding the ‘V substrate’ using the AOT. The cartoons at the top show the experimental configuration, similar to that of the ‘O substrate’, except that the end separations at the bottom end can be substantial. The ‘V substrate’ is held under 2 pN and torsionally constrained between the bottom surface of an optically trapped quartz cylinder and the coverslip surface of a sample chamber, creating a configuration with a small end separation at one end and a variable separation at the other end. The bottom plots show the measured extension and torque during braiding of the ‘V substrate’ (*N* = 17 molecules) that fall into two mean torque gap (*τ*_gap_) values, along with a direct comparison with the MC simulation results (dots) with *b* chosen to achieve reasonable agreement with measurements. For *τ*_gap_ = 3.4 pN·nm, simulations are performed with *b* = 421 nm. For *τ*_gap_ = 5.5 pN·nm, simulations are performed with *b* = 850 nm. Source data are provided as a Source Data file. **b.** Torsional measurements of braiding the ‘U substrate’ using the AOT. The cartoons at the top show the experimental configuration, similar to that of the ‘O substrate’ and ‘V substrate, except that the end separations at both ends can vary substantially. The two plots on the left show the measured extension and torque during braiding of the ‘U substrate’ (*N* = 18 molecules) that fall into two mean torque gap values, 4.8 and 6.0 pN·nm. These plots are directly compared with MC simulation results (dots) with *a* and *b* chosen to achieve reasonable agreement with measurements: assuming *a* (= *b*) = 270 nm and 543 nm respectively. The two plots on the right show the measured extension and torque within the first ±0.5 turns of braiding of the ‘U substrate’ (*N* = 40 molecules) that fall into hat tip sizes: 19 nm, 38 nm, and 56 nm. These plots are directly compared with MC simulation results (dots) with *a* and *b* chosen to achieve reasonable agreement with measurements: *a* (= *b*) = 288 nm, 408 nm, and 500 nm, respectively. Source data are provided as a Source Data file.

The ‘U substrate’ has additional striking features within ±0.5 turns (Fig. 3b). We detected two torque overshoots: a positive overshoot centered around +0.25 turns and a negative overshoot centered around -0.25 turns. These overshoots can be one order of magnitude greater than the torque gap. They are detected at similar amplitudes when turns are added to the braids in either direction (Supplementary Fig. 4). Because the overshoots peak sharply above the smoother torque profile that occurs after braiding initiates, we characterize their amplitude using the parameter ‘torque overshoot’ *τ*_overshoot_. Concurrent with observed ‘torque-overshoots’ are sharp extension drops from 0 to ±0.5 turns when the two ‘daughter’ DNA molecules are first brought in contact to initiate the braiding. We refer to this extension drop as the ‘hat tip’ ℎ_hat_tip_, because the extension-turns relation is sometimes called a hat curve.

To evaluate the geometries for the V substrate and the U substrate, we performed MC simulations with various combinations of *a* and *b* and identified those that best match the measurements. These simulation results indicate that the V substrate typically has a significant separation at only one end, while the U substrate has significant separations at both ends, consistent with the expected geometries of these templates (Supplementary Fig. 1b,c). We will follow up in the section below with a more rigorous comparison of the measurements and simulations and provide detailed interpretations.

These torsional features of the ‘V substrate’ and ‘U substrate’ demonstrate that the geometry of the braiding substrates significantly alters the torsional mechanics of braiding. The significant torque gap represents a ‘torsional barrier’ to the initiation of braiding. The dramatic torque overshoot leads to a non-monotonic torque dependence on the DNA superhelical density, creating a ‘kinetic torsional barrier’ to braiding (Supplementary Fig. 5). Thus, even if braiding is thermodynamically favored at a given torque, its onset can be delayed by the high barrier to initiation.

### Modeling How Braiding Geometry Alters Torsional Mechanics

The torsional measurements from the ‘V substrate’ and ‘U substrate’ in Fig. 3 show that as the end separations increase, several features emerge and become significant. Most notable features include a torque overshoot, hat tip, torque gap, and hat-top slope, which are further clarified in Fig. 4a. Understanding how these torsional features depend on the braiding geometry requires theoretical guidance. Thus, we performed MC simulations to further explore the torsional properties of DNA braiding over a broader range of combinations of end separations *a* and *b*. For completeness, we also modified a previous geometric model ^28^ of braiding to account for braiding geometry with different separations at the two ends (Methods; Supplementary Table 2).

**Fig. 4.**
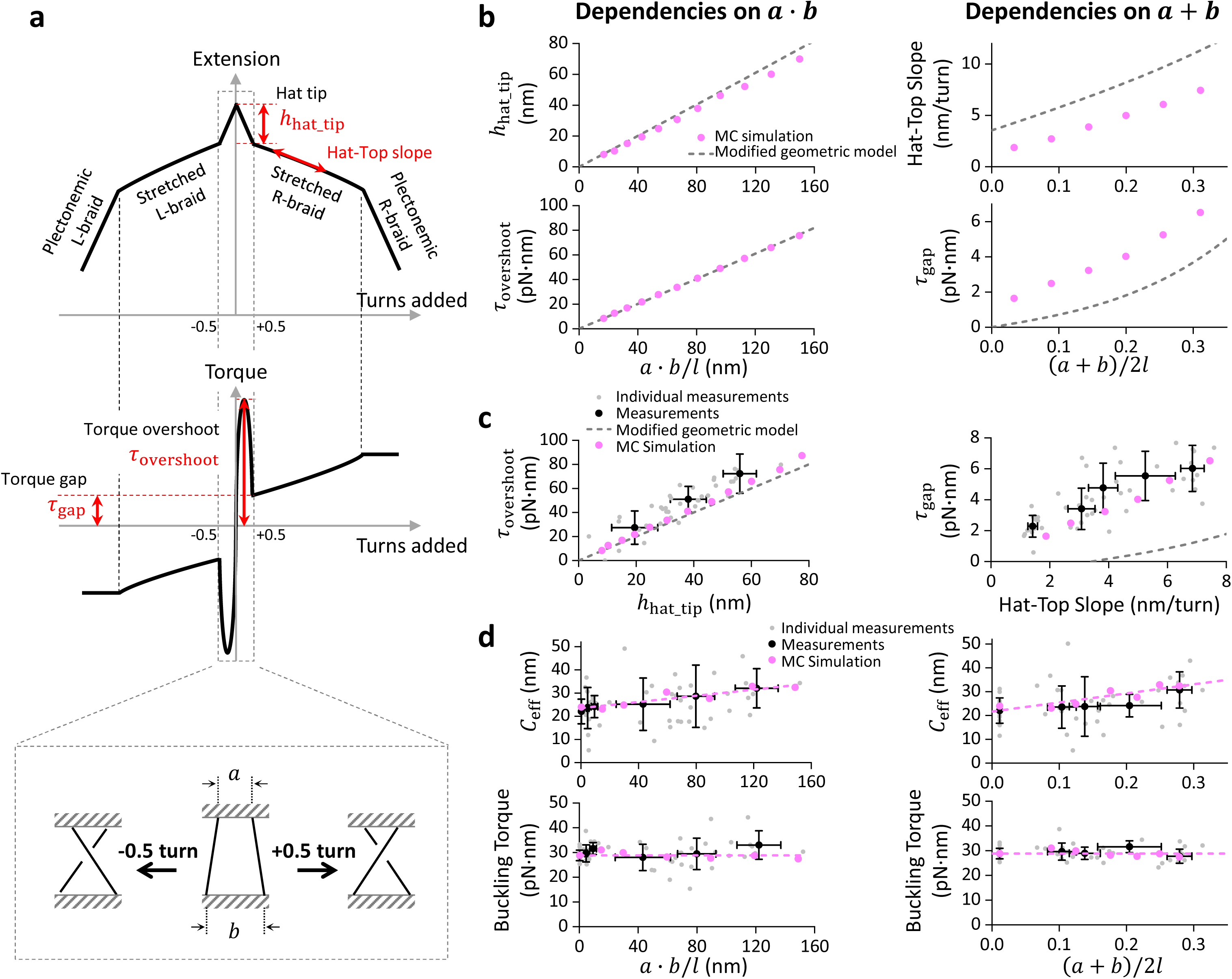
Braiding geometry governs the torsional properties of DNA braiding. **a.** A sketch depicting torsional responses of a double-DNA held at a constant force. We consider a general condition in which both *a* and *b* are substantial. From 0 to ±0.5 turns (cartoons shown at the bottom), the extension-turns relation (also called the hat curve) shows a sharp decrease in extension. Concurrently, the torque-turns relation shows dramatic, sinusoidal-like torque spikes, peaking at ±0.25 turns. We defined the extension drop as ℎ_hat_tip_ and the amplitude of the sinusoidal-like torque profile as *τ*_overshoot_. At ±0.5 turns, the two DNA strands are first brought into contact with each other, initiating braiding with a nonzero torque, which we defined as *τ*_gap_. As more turns are introduced, the two DNA strands intertwine, leading to a near-linear extension slope, which we refer to as the hat-top slope, followed by the buckling of braided DNA and formation of a plectoneme. Source data are provided as a Source Data file. **b.** MC simulations of dependencies of ℎ_hat_tip_, *τ*_overshoot_, hat-top slope magnitude, and *τ*_gap_ on braiding geometry at 2 pN. The two plots on the left show how ℎ_hat_tip_ and *τ*_overshoot_ depend on *a* · *b*, while the two plots on the right show how the hat-top slope magnitude and *τ*_gap_ depend on *a* + *b*. For comparison, the predictions from the modified geometric model (Methods) are also shown. Source data are provided as a Source Data file. **c.** Experimental validation of the MC simulations shown in **b**. To directly compare the simulation results with measurements, we plot the measured *τ*_overshoot_ versus ℎ_hat_tip_ and the measured *τ*_gap_ versus the hat-top slope magnitude using data from Fig. 2 and Fig. 3. The data points from individual molecules are shown as a scatter plot (grey dots) and are further grouped and shown as mean ± SD. For the grouped data points of the *τ*_overshoot_ versus ℎ_hat_tip_ plot, *N* = 13, 18, and 9 individual molecules from small to large *τ*_overshoot_. For the grouped data points of the *τ*_gap_ versus the hat-top slope magnitude plot, *N* = 15, 12, 7, 5, and 11 individual molecules from small to large *τ*_gap_. Source data are provided as a Source Data file. **d.** Dependencies of the effective twist persistence length *C*_eff_ and buckling torque on braiding geometry from both measurements and simulations. The data points from individual molecules are shown as a scatter plot (grey dots) and are further grouped and shown as mean ± SD. The left two plots show how *C*_eff_ and the buckling torque depend on *a* · *b*. For the grouped data points, *N* = 15, 12, 5, 10, 15, and 6 individual molecules from small to large *a* ⋅ *b*. The right two plots show how *C*_eff_ and the buckling torque depend on *a* + *b*. For the grouped data points, *N* = 15, 12, 7, 5, and 11 individual molecules from small to large *a* + *b*. The dashed lines are linear fits to the simulation results to guide the eye. Source data are provided as a Source Data file.

Our simulations show that the hat tip and the torque overshoot, i.e., torsional properties within ±0.5 turns, are essentially proportional to the product *a* · *b* (Fig. 4b, left; Supplementary Figs. 6a, 7a, and 8a). To compare simulations with measurements, we examined the torque overshoot versus hat tip and found good agreement (Fig. 4c, left). These results reveal that a torque overshoot will occur only if both ends of a braiding substrate have a substantial separation. Consistent with this conclusion, we did not detect any torque overshoot when braiding the ‘O substrate’ (Fig. 2a) or the ‘V substrate’ (Fig. 3a), which have a minimal end separation at one end or both ends. In contrast, the ‘U substrate’ can have substantial end separations at both ends and readily yield a detectable torque overshoot (Fig. 3b). In addition, predictions from the modified geometric model are in reasonable agreement with both the simulations and measurements, indicating that this simple model can be used to interpret the torsional properties of braiding within ±0.5 turns before braiding initiates.

Our simulations also show that the hat-top slope magnitude and the torque gap, i.e., torsional properties at or beyond ±0.5 turns, increase monotonically with *a* + *b* (Fig. 4b, right; Supplementary Figs. 6b, 7b, and 8b). We again compared simulations and measurements by examining the torque gap versus the hat-top slope magnitude and found them in good agreement (Fig. 4c, right). These results show that a torque gap will appear if at least one end of a braiding substrate has a substantial separation. Indeed, we detected a distinct torque gap when braiding either the ‘V substrate’ (Fig. 3a) or the ‘U substrate’ (Fig. 3b), while the torque gap of braiding the ‘O substrate’ was minimal (Fig. 2a). Importantly, predictions from the modified geometric model deviate significantly from both simulations and measurements (Supplementary Fig. 9). While this simple model works well before braiding formation, it neither accurately describes torsional properties once braiding begins nor predicts braid buckling. Thus, the geometric model cannot provide guidance on how braiding geometry affects the torque gap, the effective torsional modulus of a braid, and the braid buckling. Therefore, we will not use it further in subsequent analysis. This highlights the need for simulations to understand the geometry-governed torsional properties of braiding.

We further examined how braiding geometry modulates the effective twist persistence length *C*_eff_ and the buckling torque by comparing simulations and measurements (Fig. 4d). To characterize the braiding geometry in the measurements, we estimated *a* · *b* and *a* + *b* from the measured hat tip size and the measured hat-top slope, respectively, using the simulated relations shown in Fig. 4b. We first examined how *C*_eff_ depends on these geometric parameters. Both simulations and measurements show little dependence of *C*_eff_ on *a* · *b* or *a* + *b* (Fig. 4d). We then examined how the buckling torque depends on these geometric parameters. Both simulations and measurements show that the buckling torque remains essentially unchanged with either *a* · *b* or *a* + *b* (Fig. 4d). Thus, plectoneme formation of braided DNA occurs at a critical torque, regardless of the substrate geometry. Consequently, when a large torque gap is present, the turns required to reach the critical buckling torque are reduced (Fig. 3; Supplementary Fig. 2b,c).

Thus, both measurements and simulations show how the braiding geometry modulates the torsional mechanics of braiding. These results provide an explanation for how large end separations lead to larger torque gaps and torque overshoots, thereby creating a ‘kinetic torsional barrier’ to the initiation of braiding.

## Discussion

Our work demonstrates that DNA braiding geometry plays a crucial role in the torsional mechanics of DNA braiding. These findings have significant implications for DNA supercoiling partitioning and the torsional resistance experienced by the replisome. During replication, DNA supercoiling generated by replication must be distributed either ahead of or behind the replisome. Supercoiling partitioning behind the replisome will lead to fork rotation and intertwine (braid) the two daughter DNA molecules and must be untangled by topoisomerases before chromosome segregation. Our work allows prediction of this supercoiling partitioning and the accompanying torsional stress.

For simplicity, we consider a case where DNA replication takes place over a naked DNA and proceeds to the middle of the DNA substrate. Since the daughter strands are brought together at one end by the replisome and at the other end by SMC proteins or another replisome, the braiding geometry resembles the ‘O substrate’ (Fig. 5a). If topoisomerase activity nearly keeps up with replication, the DNA superhelical density is low. Since the torsional stiffness of braided DNA is much smaller than that of a single DNA molecule, replication will lead to fork rotation, and the supercoiling will partition predominantly behind the replisome. Thus, although the small end separations help alleviate some torsional stress, they will also lead to daughter DNA intertwining. If topoisomerase activity cannot keep up with replication, the DNA superhelical density may increase, leading to buckling of the parental single-DNA strand at the front and limiting further accumulation of torsional stress. Concurrently, the fork stops rotating, and additional supercoiling will partition to the front of the replisome.

**Fig. 5.**
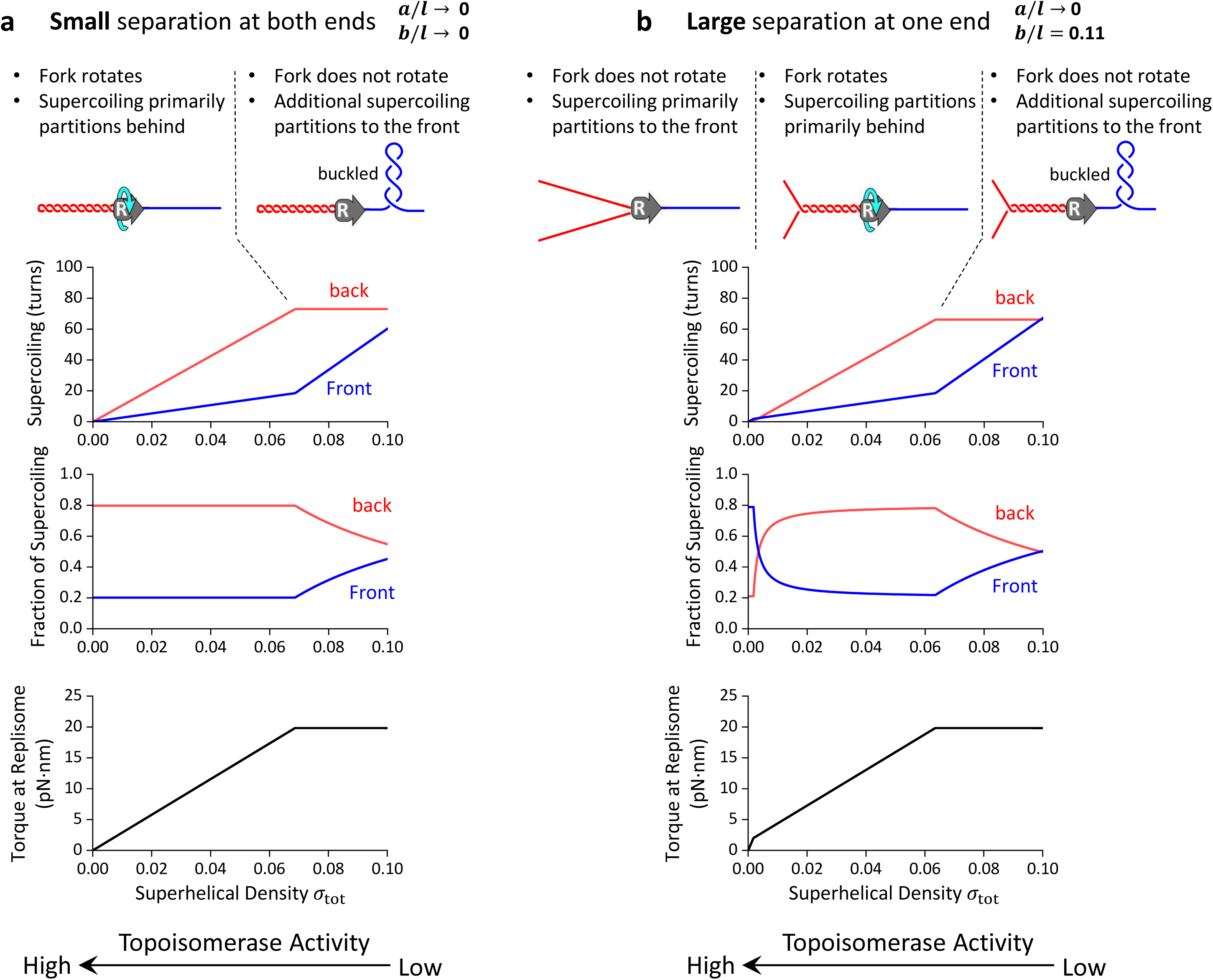
Supercoiling partitioning during DNA replication altered by different substrate geometries. Shown are predictions of two example substrate geometries when the replisome has replicated to the middle of the DNA template (a total length of 14 kb). The predictions are based on the torsional mechanical measurements of a single DNA-molecule substrate (in front of the replisome) and a double-DNA molecule substrate (behind the replisome). For simplicity, the illustrative plots are obtained by considering only the linear term of torsional stiffness of the substrate at 2 pN (Supplementary Table 3). **a.** Daughter-strand geometry with small separation at both ends: *a*/*l* = *b*/*l* ≍ 0. Under this geometrical configuration, there is no torsional barrier to limit supercoiling partitioning behind the replisome. If topoisomerase activity is high, the superhelical density is low. Under this condition, significant fork rotation can occur. The intertwining of the daughter strands prevents chromosome segregation but buffers torsional resistance to replisome progression. If topoisomerase activity is reduced, the superhelical density increases, and so does the torque at the replisome. This eventually leads to the single-DNA substrate at the front buckling to form a plectoneme, which is highly effective at limiting torsional stress accumulation. Once buckling occurs, the fork stops rotating, leading to additional supercoiling partitioning only to the front. **b.** Daughter-strand geometry with a large separation distal to the replisome: *a*/*l* ≍ 0 and *b*/*l* = 0.11. This may occur when SMC proteins fail to bring the two daughter strands together at the end distal to the replisome. Under this geometrical configuration, braiding the daughter strands must first overcome a ‘torsional barrier’ presented by the torque gap that resists the initiation of braiding and limits daughter-strand intertwining. If the replisome overcomes this barrier, the fork rotates, and supercoiling partitions preferentially behind the replisome. This continues until the single-DNA substrate in the front buckles, limiting the accumulation of torsional stress. Once buckling occurs, the fork stops rotation, and additional supercoiling will be partitioned only to the front of the replisome.

In contrast, if the daughter strands are brought together at one end by the replisome but fail to be brought together at the other end by SMC proteins, then the braiding geometry resembles the ‘V substrate’ (Fig. 5b). This geometry will result in a torque gap, which imposes a ‘torsional barrier’ to braiding initiation. If topoisomerase activity is high and can nearly keep up with replication, the DNA superhelical density will be low. The fork will not rotate due to the presence of the ‘torsional barrier’ imposed by the torque gap against braiding the daughter DNA strands, and thus supercoiling will be directed to the front of the replisome. Although the two daughter DNA molecules are minimally intertwined, the replisome experiences a greater torsional stress. Interestingly, previous studies show that DNA intertwining is reduced upon inactivation of cohesin ^71^ and the progression of the replication fork is slowed in cohesin-deficient cells ^72^, providing possible evidence for our proposed model. If topoisomerase activity cannot keep up with replication, an increase in the superhelical density will braid the daughter strands and subsequently buckle the single DNA substrate in the front. Once buckling occurs, torque ceases to increase, and the fork stops rotating. Any additional supercoiling will partition to the front of the replisome.

The framework presented here captures key features of supercoiling partitioning and suggests that geometry is a major determinant of this process. While these conclusions are grounded in experimental and theoretical results for naked DNA, their broader applicability will depend on understanding how additional factors, particularly bound proteins, modify torsional mechanics. For instance, converting naked DNA to chromatin has been shown to significantly change the torsional mechanics of the substrates ^4^, in turn affecting supercoiling partitioning. How the geometry of protein-bound braiding substrates influences supercoiling partitioning remains to be explored, offering a valuable avenue for future investigation. Extending this work to physiologically relevant fork substrates, rather than separate examinations of single- and double-DNA substrates, would provide a more integrated view of partitioning. Moreover, because torque measurements are central to this analysis, enhanced torque resolution, such as that enabled by smaller cylinders in the AOT ^55^ or by the rotary bead assays ^73,74^, may reveal subtler torque gaps and overshoots and more precisely characterize the torsional response observed during the first half turn. Together, these advances would enable a more complete and physiologically relevant understanding of supercoiling partitioning during replication across diverse substrate geometries and protein-bound contexts.

## Methods

### Plasmid Cloning

The braiding templates ‘O substrate’ and ‘V substrate’, were created from pMDW168. This plasmid was constructed via Gibson assembly and site directed mutagenesis. The Gibson assembly used four dsDNA segments: two ∼ 7 kb segments for the arms of the braiding substrate and two ∼ 70 bp segments for the anchors. Both ∼ 7 kb segments were amplified via PCR from pMDW133 using primers (Arm1 F, Arm1 R, Arm2 F, Arm2 R, Supplementary Table 4) and Q5 DNAP (NEB, M0491S), followed by purification with Purelink PCR columns (Invitrogen, K310001). The ∼ 70 bp anchor segments were produced by amplifying two 60 nt ssDNA (Anchor A template, Anchor T template, Supplementary Table 4) oligonucleotide via PCR with primers (Anchor A_F, Anchor A_R, Anchor T_F, Anchor T_R, Supplementary Table 4). The resulting two anchors and two arms were mixed in a 5:1 molar ratio and assembled using NEBuilder HiFi DNA assembly mastermix (NEB, E2621S). The assembled DNA was transformed into NEB 5α cells (NEB, C2987H) to make an intermediate plasmid. This intermediate plasmid had an unwanted Nt.BbvCI (NEB, R0632) recognition sequence on top of four that was included by design. This site was removed using site directed mutagenesis, first using PCR amplification from the intermediate plasmid with primers (KLD nickRemoval F, KLD nickRemoval R, Supplementary Table 4), followed by a KLD reaction on the PCR product with T4 PNK (NEB, M0201S), T4 ligase (NEB, M0202S) and DpnI (NEB, R0176S). We transformed the final product into NEB 5α competent cells, resulting in plasmid pMDW168.

### DNA Template Construction

The preparation of ‘O substrate’ is shown in Fig. 1a. The plasmid pMDW168 was digested with Nt.BbvCI to introduce two pairs of nicks, each pair containing two nicks spaced ∼ 70 bp apart, for a total of four nicks on the plasmid. Two ∼ 70-nt ssDNA gaps were then created by incubating the nicked plasmid with excess competitor ssDNA with complementary sequence (Dig-end competitor 1 and 2, Bio-end competitor 1 and 2, Supplementary Table 4) using a thermocycler and the following program: 15 min at 82 °C, then 42 cycles of 1 min with -1 °C temperature gradient per cycle. The two created ssDNA gaps were filled with digoxigenin (Roche, 11093088910) and biotin (Invitrogen, 19524016) labeled nucleotides via Hemo KlenTaq (NEB, M0332). The Hemo KlenTaq does not have strand displacement or exonuclease activity, resulting in a filled gap with an unligated nick at the end. The final product was purified with Purelink PCR purification columns. The resulting circular DNA had two torsionally unconstrained dsDNA segments free for braiding, with each end separation constrained by ∼ 70 bp.

The preparation of ‘V substrate’ template is shown in Fig. 1b. The plasmid pMDW168 was linearized by digestion with StyI-HF (NEB, R3500). The ∼ 500 bp long biotin labeled adapter was amplified via PCR from the plasmid pMDW111 with ∼ 25% of the dATP replaced with biotin-dATP. The PCR was performed using Taq DNAP (NEB, M0273) and primers (0.5 kb PCR F, 0.5 kb PCR R, Supplementary Table 4). The amplified adapter DNA was digested with StyI-HF to produce overhangs matching the linear template. The two digested products were ligated together with T4 ligase, and then purified with Purelink PCR purification columns. The product was nicked with Nb. BbvCI (NEB, R0631) and incubated with the competitor ssDNA (Dig-end competitor flip 1 and 2, Supplementary Table 4) to create one 70-nt ssDNA gap using the following program: 15 min at 82 °C, then 42 cycles of 1 min with -1 °C temperature gradient per cycle. The ssDNA gap was then filled with digoxigenin labeled nucleotides via Hemo KlenTaq. The final product was purified with Purelink PCR purification columns. The resulting ‘V substrate’ has one DNA segment torsionally constrained and the other one torsionally unconstrained.

The 6.5 kb torsionally unconstrained template was amplified via PCR from plasmid pMDW133, with primers (6.5 kb PCR F, 6.5 kb PCR R, Supplementary Table 4) ^57^.

The 12.7 kb torsionally constrained template was first amplified via PCR from λ DNA, with primers (12.7 kb PCR F, 12.7 kb PCR R, Supplementary Table 4). The products were digested with AvaI (NEB, R0152) and BssSI-v2 (NEB, R0680), then ligated with multi-labeled adapters as described previously ^58^.

### Angular Optical Trapping

An angular optical trap (AOT) is ideally suited for DNA torsional studies because it permits simultaneous measurements of force, position, torque, and rotation of a trapped quartz cylinder ^75,76^. The quar{Citation}tz cylinders were fabricated out of a quartz wafer (Precision Micro Optics, PSQB-131332) ^4,5,56–61,77^ based on deep-ultraviolet-lithography-based nanofabrication process ^4,57–59^. These functionalized cylinders were covalently attached with anti-digoxigenin (Roche, 11333089001) via glutaraldehyde as a linker (Sigma, G5882) ^77^. These cylinders have dimensions similar to those previously fabricated ^57^, with a height of about 1 µm and a diameter of about 0.5 µm. Since only the end of each cylinder is functionalized for DNA attachment, the diameter of the cylinder places an upper limit on the end separation *a* for the ‘U substrate’.

### Single-Molecule Chamber Preparation

Single-molecule sample chambers for DNA torsional studies using the AOT ^78^ were assembled with a nitrocellulose-coated coverslip (Fisher Scientific, 12544B) and a Windex-cleaned glass slide (Fisher Scientific, 1255010) ^4,59^. The chamber surface was functionalized with streptavidin (Agilent, SA10) conjugated to a coverslip coated with biotinylated bovine serum albumin (ThermoFisher, 29130) ^77^. The functionalized chamber was passivated with β-casein (Sigma, C6905) prior to tethering DNA molecules.

For both the ‘O substrate’ and the ‘V substrate’, the prepared braiding substrate was immobilized on the functionalized coverslip surface. After flushing out free DNA within the chamber, the anti-digoxigenin coated quartz cylinders were flowed in to attach to the surface-immobilized DNA. In addition, the ‘V substrate’ has multiple Nt. BsmAI nicking sites on both strands for braiding (see pMDW168 sequence below for details) and was nicked with 30 U/mL Nt. BsmAI (NEB, R0121) in the nicking buffer (10 mM Tris pH 8.0, 10 mM NaCl, 0.5 mM MgCl_2_, 1.5 mg/mL β-casein) for 10 min after the cylinder attachment step to prevent torsion accumulation within individual ‘daughter’ DNAs. Previously, we systematically studied the efficiency of nicking enzymes and found that 30 U/mL Nt. BsmAI for 10 min nicked 100% of a template containing 9 nicking sites (see Extended Data Figure 5c of reference ^79^). In this work, we also used 30 U/mL Nt. BsmAI for 10 min to nick a DNA template containing 12 nicking sites. This template should be fully nicked.

For the ‘U substrate’ braiding assay using the 6.5 kb DNA, the DNA was first mixed with the quartz cylinders to increase the fraction of double-DNA tethers and then introduced to the sample chamber for surface immobilization.

Prior to torsional measurements on the AOT, each chamber was flushed with the topoisomerase reaction buffer (10 mM Tris-HCl pH 8.0, 50 mM KCl, 50 mM NaCl, 3 mM MgCl_2_, 1 mM ATP, 0.1 mM EDTA, and 1.5 mg/mL β-casein) ^59,80^. Unless otherwise stated, all these processes were performed at room temperature (23 °C).

### Data Acquisition and Analysis

The torsional measurements for braided DNA molecules using the AOT were performed at a laser power of 40 mW entering the objective to minimize the possibility of photo-damage while preventing cylinder slippage ^55,81^. The experimental procedures were similar to those for a single-DNA molecule ^56–59,62–64,82,83^. In each measurement, the tethered quartz cylinder was held at a constant force and rotated at 2 turn/s to add or remove turns to the braiding substrate. Extension and torque were simultaneously measured at 10 kHz and averaged to 400 Hz. The measured extension had a slight but non-negligible offset. To more accurately determine the extension, we corrected this offset using the extension predicted by the worm-like-chain model at zero turns ^84^. Torque data was filtered with a 2 s sliding window to reduce the level of Brownian noise. To resolve the torque overshoot that occurs within ±0.5 turns (Fig. 3b, right plots), the torque filter sliding window size was reduced to 0.01 s near the zero-turn state. For comparison, we performed the torsional measurements and analyzed the data for single-DNA substrates under the same conditions. All the measurements were performed at room temperature (23 °C). To reduce Brownian noise, multiple traces were averaged to characterize the braiding substrates. Since the end separations of the ‘V substrate’ and the ‘U substrate’ varied, traces in Fig. 3a and left plots of Fig. 3b were grouped by the torque gap, while traces in the right plots of Fig. 3b were grouped by the hat tip. Such grouping was performed to reduce the Brownian noise in torque measurements, though it somewhat obscured the buckling transition in the averaged trace (as the geometries of ‘V substrate’ and ‘U substrate’ were not fully controlled).

The effective twist persistence length 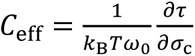 of braided DNA was determined by linearly fitting the torque slope within 5/*Lk*_0_ < *σ*_c_ < 0.05 for individual torsional measurements, where *τ* is the measured torque, *σ*_c_ is the catenation density, *Lk*_0_ is the intrinsic linking number of each ‘daughter’ DNA molecule, *k*_B_*T* the thermal energy, and 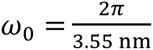. The buckling transition of each braided DNA was determined by fitting the extension-turns relation with a 2-piecewise function, which consists of a quadratic region for the pre-buckling regime and a linear region for the post-buckling regime. The buckling torque was determined by averaging the torque data after the buckling transition, as the post-buckling torque plateaus under our experimental condition (Fig. 2a; Fig. 3a; Fig. 3b). The hat tip size ℎ_hat_tip_was determined by fitting the individual hat curves within ±0.5 turns with equation (7). The hat-top slope was obtained by linearly fitting the extension slope within 5/*Lk*_0_ < *σ*_c_ < 0.05 of individual hat curves. The *τ*_gap_ was obtained from half the torque difference between the two linearly fitted torque curves around the zero turns.

#### Coarse-Grained Monte Carlo simulations of DNA Braiding

We modeled DNA molecules as discrete worm-like chains and simulated equilibrium configurations of different braiding geometries using a Monte Carlo (MC) procedure. We extended a previous simulation framework ^28^ by incorporating a stronger topological invariant to constrain topology and a Debye-Hückel interaction potential ^28,70^ as detailed below. We simulated equilibrium conformations for a braided template with top separation *a* and bottom separation *b* for a range of turns added (or catenations) *n* held under a constant force *F* applied along **Ẑ**. The thermodynamic potential for such a template consists of the following terms:

1. Bending energy of the two DNA molecules:

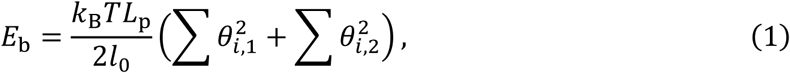

where *θ_i_*_,*X*_ is the angle between two consecutive segments *i* and *i* + 1 in chain *X*, *k*_B_*T* is the thermal energy, *L*_p_ is the linear persistence length and *l*_0_ is the discrete segment length. To mimic the anchor design of the ‘O substrate’, we also included bending penalties at the four anchor points of the two DNA chains. Similarly, for the ‘V substrate’, bending penalties at the top anchor points were included in the potential.

2. Electrostatic interaction potential modeled using an approximate Debye-Huckel potential:

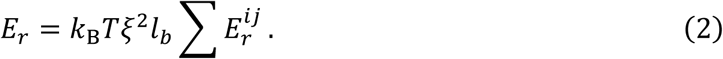

Here, *ξ* is the effective charge density, *l_b_* is the Bjerrum length in water at the temperature *T*, and 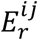 is solved numerically by computing 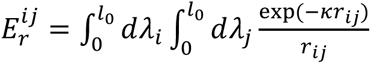 for all pairs of non-adjacent segments *i* and *j*. Here, the integration is done along the two segments, *λ_i_* and *λ_j_* are the DNA contour parameters, *r_ij_* is the distance between the two positions on the segments corresponding to *λ_i_* and *λ_j_*, and finally, *κ* is the inverse of the Debye length (Supplementary Table 1). A look-up table for the double integral was created for a range of 3D segment orientations as previously described ^85,86^.

3. Spring potentials that constrain the distance 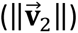 and direction (**V̂**_2_) of the top-anchor points of the two DNA chains, akin to those implemented in ^28^:

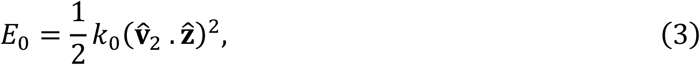

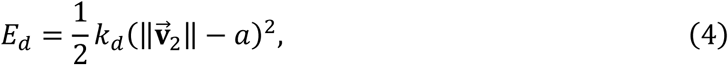

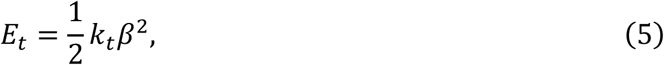

with *k*_0_, *k_d_*, *k_t_* are fictitious spring constants (Supplementary Table 1) and *β* is the angle deviation of top separation vector **V̂**_2_ from desired vector direction **â**. The bottom DNA anchors and the corresponding separation *b* remain unchanged throughout the course of the simulation, and hence, do not need to be explicitly constrained by spring potentials. Note that *E_t_* constrains angular fluctuations and functions as a torsional spring detector that allows us to obtain the torque, using *τ* = *k_t_β*.

4. Finally, an additional term:

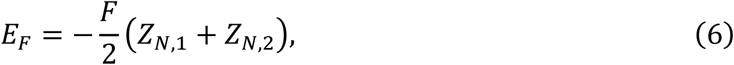

is added, given that the thermodynamic potential considered is under a constant force. *Z_N_*_,*X*_ is the Z-position of the final bead of chain *X*.

The parameters used in this work to compute the energy terms are given in Supplementary Table 1. Here, we provide details about how the parameters are chosen. The value of *L*_p_ was chosen to be comparable to the persistence length measured in our experimental buffer. A short segment length of *l*_0_ = 5 nm was then selected to be much smaller than *L*_p_, allowing for smooth curvatures in the DNA backbone. For the value of spring constants, we balanced the need for stiff springs that effectively constrain anchor geometry and orientation as desired, while allowing sufficient thermal fluctuations to explore different configurations for computational efficiency. We found the combination of spring values shown in Supplementary Table 1 to work well for our system. For the Debye-Hückel parameter, we explored a range of parameter values to find a set of *κ* and *ξ* that effectively captured the measured torsional properties. Since our experimental buffer contains divalent cations (with chelating agent) in addition to monovalent salt, direct estimations of *κ* and *ξ* were not feasible. The combination of *κ* and *ξ* in Supplementary Table 1 produced a reasonable match for all different geometries and forces shown in the manuscript.

We began the simulations by preparing a pair of discrete helices intertwined with the desired value of catenation (*n*) as initial conformations. We then sample the conformational landscape using three different classical polymer moves for each of the two chains separately: pivot ^87^, crankshaft ^88^, and reptation ^88^. A new braided confirmation was accepted after a two-stage check. The first stage was the Metropolis-Hasting condition that ensured convergence to equilibrium. The second stage ensured that the topology remains unchanged throughout the course of the simulation, by computing the bivariate Alexander polynomial numerically at values of -1.1 for each of the two variables. Previous studies reported that substituting each of the variables with -1 was efficient for constraining topology ^70^, however we found that it was not sufficient, due to degeneracy in topology results. Introducing a rigorous topology check allowed us to simulate torsional mechanics into the buckling regime for different configurations, which has previously not been reported through simulations. Finally, we tuned the range of each of our moves such that the final acceptance ratio remains between 15% to 40%.

The methodology used to simulate the torsional properties of a single-DNA substrate was described in a previous work ^58^. The procedure used for simulating single-DNA twisting shares many details with the braiding procedure described above. Here, we briefly highlight some of the differences. First, since the single-DNA substrate is torsionally constrained (and not nicked), we added a twisting penalty to the thermodynamic potential term. The twisting penalty was indirectly calculated by invoking the conservation of linking number Δ*Lk* = Δ*Tw* + Δ*Wr*, where Δ*Lk* is the change in linking number from the relaxed value (i.e., turns added to the DNA), Δ*Tw* is change in overall twist and Δ*Wr* is the global writhe change. During the simulation, we numerically computed Δ*Wr* of the chain contour and assigned twist, Δ*Tw* = Δ*Lk* − Δ*Wr* to the rest of the chain. This allows us to directly ensure the correct linking number without requiring fictitious springs (like equation 5). The other main difference was how we ensured correct global topology for sampled conformations. For the single-DNA substrate, we used a single-component Alexander polynomial to ensure that the configuration remains unknotted during the simulation. Other details for single DNA simulations (like remaining energy terms and simulation moves) remained the same as those described for our braided template simulations above.

We validated our simulations by comparing the simulation results with the torsional measurements of the ‘O substrate’ and the single-DNA substrate (Fig. 2a,b). For the ‘O substrate’, we simulated two 2130 bp DNA chains separated by 21 bp. These dimensions are about 3 times smaller than those of the ‘O substrate’ primarily to reduce computational cost. The resulting braided simulations are then linearly scaled up to compare with measurements. Note that once braiding starts (*n* > 0.5), both extension and torque curves are only functions of (*a* + *b*)/2*l* regardless of the DNA contour length (Fig. 4; Supplementary Figs. 6, 7, and 8). Hence, linear scaling is valid in the braiding regime. After systematically simulating braiding with an extensive range of braided geometries, we also directly compared our simulated results with the torsional measurements of the ‘V substrate’ and the ‘U substrate’ (Fig. 3a,b). For both V and U substrates, we also simulated substrates about 3 times shorter than those used in experiments to reduce computational cost and scaled the results linearly in the braiding regime (*n* > 0.5) when comparing with experiments. For the U substrate in the (*n* < 0.5) regime, since hat-curve and torque are functions of (*a* · *b*)/*l*, we simulated the non-scaled DNA lengths 6564 bp before comparing against experiments (Fig 3b, right). Two snapshot configurations of each substrate, one before buckling and one after buckling, are shown in Supplementary Fig. 10. Finally, the simulations are utilized to infer braiding end separations *a* and *b* of our measurements and understand their impact on the torsional mechanics of DNA braiding (Fig. 4).

We also used our simulations to estimate the ‘effective’ separation at the multi-tag anchors engineered in ‘O substrate’ and ‘V substrate’. While the end separation is well defined by the multi-tag sequence length (∼70 bp or ∼24 nm contour length), the effective separation between the two strands is expected to be greater than 24 nm since the bending curvature cannot have discontinuities at the anchors due to the finite DNA persistent length. To estimate this effective separation, we compared the simulation results for the ‘O substrate’ and ‘U substrate’ with *a* = *b* = 24 nm, both with no turns added. Simulations of the ‘O substrate’ required consideration of the DNA bending penalty at the anchors, whereas those of the ‘U substrate’ has no bending penalties considered at the anchors. We analyzed the equilibrated configurations for both substrates to estimate the average Euclidean separation distance between the two chains at two vertices ∼1 Kuhn length away (∼100 nm) along the DNA contour from each of the two anchor points. The difference between these separation distances for ‘O substrate’ and the ‘U substrate’ gives an upper bound estimate for the extra separation required to bend the DNA at the anchor points of the ‘O substrate’. Using this analysis, we estimate an effective end separation of the ‘O substrate’ to be 47, 46, 42, and 41 nm at 0.5, 1.0, 2.0, and 3.0 pN, respectively.

### Modified Geometric Model of DNA Braiding

In addition to the MC simulations, we also modified a previous geometric model ^28^ of braiding DNA to consider the general case with different separations at the two ends. In this model, each DNA is assumed to be a bendable rod with a fixed length calculated from the end-to-end extension of the DNA under stretching forces at zero turns, and the braiding angle *α* is assumed to be the same across all the braiding regions. We obtained the following expressions for the extension and torque, under a given force (*F*) and number of turns added (*n*).

The extension *z* of the braiding substrate is

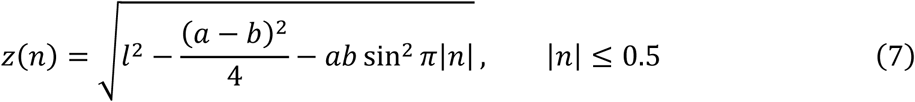

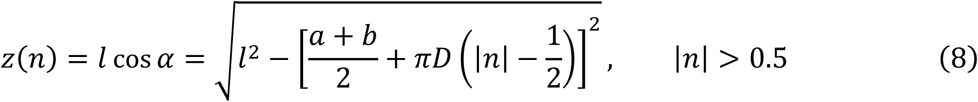

where *n* is the catenation number, *l* is the length of the bendable DNA rod, *a* and *b* are the end separations, and *D* is the diameter of a DNA braid.

When 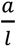 and 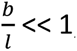, the size of the hat tip (Fig. 4a) from equation (7) is

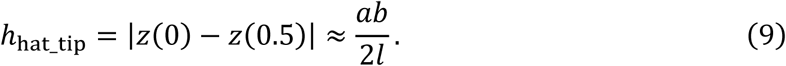

Beyond 0.5 turns, the modified geometric model gives a slightly non-linear extension-turn relationship with a catenation-number-dependent hat-top slope:

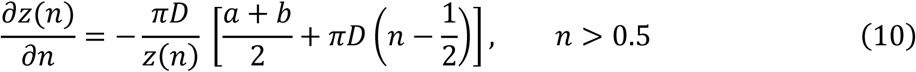

This non-linearity makes it difficult to directly compare the hat-top slope from the geometric model with those from simulation and experiments, since the latter two show a more linear relationship. To facilitate the comparison, we imposed a linear fit on the extension-turn relationship generated by the modified geometric model between 0.5 turns and *σ*_c_ < 0.05. This fitted slope was then defined as the hat-top slope from the modified geometric model (Fig. 4a).

As turns are introduced, braiding must work against the stretching force to shorten the extension *z*(*n*), requiring energy:

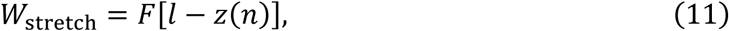

and overcome the DNA bending energy:

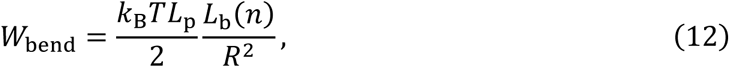

where *F* is the axial stretching force, *L*_p_ is the DNA linear persistence length, *L*_b_(*n*) is the total DNA contour length in the braid, and *R* is the curvature radius of the braided helix. The total energy introduced into the braiding substrate is *E*(*n*) = *W*_stretch_(*n*) + *W*_bend_(*n*). The braiding torque is thus

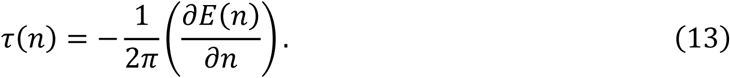

Analytically, the braiding torque is:

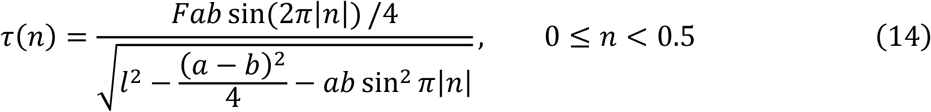

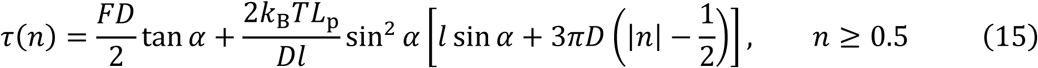

where 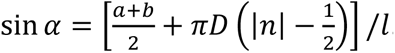. For the regime of *n* < 0, since the geometric model assumes no chirality of braiding, the torque values should have the same expressions as equation (14) and equation (15) except with a negative sign.

We obtained *τ*_overshoot_ (Fig. 4a) as the torque value at *n* = 0.25 turns and *τ*_gap_ (Fig. 4a) as the torque values at *n* = 0.5 turns. The *τ*_overshoot_ expression under the limit *a* ≪ *l* and *b* ≪ *l*:

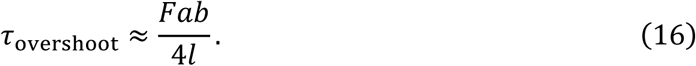

The *τ*_gap_ expression is

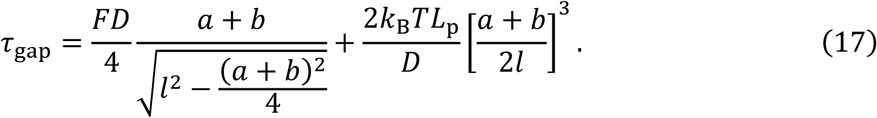

The parameters in this modified geometric model are shown in Supplementary Table 2. This model provides good estimations of ℎ_hat_tip_ and *τ*_overshoot_, while it vastly overestimates the hat-top slope magnitude and underestimates the *τ*_gap_ compared with our MC simulations and measurements for (Fig. 4b; Fig. 4c), indicating it should not be used after braiding initiates. This limitation has been previously reported ^28^.

### Calculation of Supercoiling Partition during Replication

DNA replication introduces (+) supercoiling in the DNA substrate as the parental DNA is separated to produce two daughter strands. This supercoiling must partition to either the front of the replisome (the parental DNA) or behind the replisome (daughter strands). Thus, the overall supercoiling density of a replicating DNA substrate can be written as 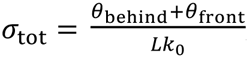, where *θ*_behind_ is the number of turns absorbed behind, *θ*_front_ is the number of turns absorbed to the front, and *Lk*_0_ is the total intrinsic linking number of the parental DNA before replication initiation.

For the illustrative plots shown in Fig. 5 and Supplementary Fig. 5, we used *Lk*_0_ = (14000/10.5) turns and 2 pN force. Considering the scenario when the replisome proceeds to the midpoint of a DNA substrate, we calculated the fraction of supercoiling partitioned behind the replisome 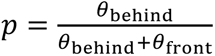 as a function of *σ*_tot_, and correspondingly, the torque at the replisome *τ*_replisome_ (Fig. 5; Supplementary Fig. 5), characterized by an effective torsional modulus of the coupled braided and single-DNA substrates ^4^. It is worth noting that the first 0.5 turns of the DNA braiding do not scale with the DNA length. For the case where *a*/*l* = *b*/*l* = 0 (Fig. 5a), the *C*_eff_ of braided DNA was directly used to characterize the torsional modulus within the first 0.5 turns. For the case where *a*/*l* = 0 and *b*/*l* = 0.11 (Fig. 5b), the torque was assumed to rise linearly to *τ*_gap_ = 2 pN·nm in the first 0.5 turns. For the case where *a*/*l* = *b*/*l* = 0.14 (Supplementary Fig. 5), the torque to initiate a DNA braid was given by equation (16). The detailed parameters used for this calculation are summarized in Supplementary Table 3.

## Data Availability

All data needed to evaluate the conclusions in the paper are present in the paper and/or the Supplementary Materials. Source data are provided with this paper. The source data file contains all data shown in the main text figures and the supplementary figures. These data are also available from the following GitHub repository link: https://github.com/WangLabCornell/Hong_et_al_2 ^89^.

## Code Availability

The code for the MC simulations and AOT data analysis can be obtained from the following GitHub repository link: https://github.com/WangLabCornell/Hong_et_al_2 ^89^.

## Acknowledgments

We thank members of the Wang Laboratory for helpful discussions, Dr. J. T. Inman for providing technical assistance in updating the AOT software, Drs. J. Lee and X. Gao for the preliminary exploration of this project, Dr. X. Jia for the helpful discussion on DNA-substrate preparation and for providing the 6.5 kb torsionally unconstrained DNA used in this work, and Dr. M. Wu for providing the 12.7 kb torsionally constrained DNA.

## Author Contributions Statement

S.P. and M.D.W conceived the design of the braiding substrates. S.P. created the original template plasmid. Y.H. and S.P. prepared the braiding substrates. Y.H. fabricated the quartz cylinders. Y.H. designed and optimized the single-molecule assays. Y.H. performed measurements with assistance from H.W.. Y.H. performed data analysis. G.S performed the MC simulations with assistance from H.W.. H.W. modified the geometric model for DNA braiding. Y.H. and M.D.W. drafted the manuscript with revisions by all authors. M.D.W. supervised the project.

