## Supplementary Information for "Geometry of Braided DNA Dictates Supercoiling Partitioning"

**Supplementary Table 1**. Parameters used in the MC simulations of DNA braiding.

| Parameter | Value |
|---|---|
| Segment length ($l_0$) | 5 nm |
| Thermal energy ($k_\mathrm{B}T$) | 4.1 pN·nm |
| DNA linear persistence length ($L_\mathrm{p}$) | 48.3 nm |
| Torsional Spring Constant ($k_t$) | 205 pN·nm/rad |
| Lateral Spring Constant ($k_d$) | 82 pN/nm |
| Vertical Spring Constant ($k_0$) | 820 pN·nm |
| Inverse Debye Length ($\kappa$) | 1.55 $nm^{-1}$ |
| Effective Linear Charge Density ($\xi$) | 5.08 $e^{-}$/nm |

**Supplementary Table 2**. Parameters used for the modified geometric model of DNA braiding.

| **Parameter** | **Value** |
|---|---|
| Thermal energy ($k_{\mathrm{B}}T$) | 4.1 pN·nm |
| Linear persistence length ($L_{\mathrm{p}}$) | 48.3 nm |
| Diameter of a DNA braid ($D$) | 6.5 nm |

**Supplementary Table 3**. Parameters used for calculating supercoiling partition during replisome elongation, related to Fig. 5 and Supplementary Figure 5.

| | $a/l = b/l \rightarrow 0$ | $a/l \rightarrow 0$, $b/l = 0.11$ | $a/l = b/l = 0.14$ |
|---|---|---|---|
| Torque overshoot ($\tau_{\mathrm{overshoot}}$) | 0 pN·nm | 0 pN·nm | 20 pN·nm |
| Torque gap ($\tau_{\mathrm{gap}}$) | NA (as no discontinuity was considered for the first 0.5 turns) | 2 pN·nm | 3 pN·nm |
| Effective twist persistence length of a DNA braid ($C_{\mathrm{eff}}$) | 25 nm | 25 nm | 25 nm |
| Effective twist persistence length of a single DNA ($C_{\mathrm{eff}}$) | 98.5 nm | 98.5 nm | 98.5 nm |
| Thermal energy ($k_{\mathrm{B}}T$) | 4.1 pN·nm | 4.1 pN·nm | 4.1 pN·nm |
| Buckling torque of the single-DNA substrate | 19.8 pN·nm | 19.8 pN·nm | 19.8 pN·nm |

**Supplementary Table 4**. Primer sequences for plasmid cloning and DNA template construction.

| **Oligo Name** | **Sequence (5' → 3')** |
|---|---|
| Arm1 F | CAATGGCCTGCGTATCTTCAA |
| Arm1 R | TCCGGAGAGTCAGCGATGTT |
| Arm2 F | TCACCATTACCGGATACAACC |
| Arm2 R | TGTTACGAAGATGGATGCCT |
| Anchor A template | AACAACAAACAGAAACAGACAAGAAACAAAGCAAAGAAACAAGAGAGA<br>AAC<br>AGAACACGA |
| Anchor T template | TCGTGTTCTTTGGTTCTTTGTCTTGTTTCCTTGTTTGTTCTTTCTTGCTTGTTT<br>GTTGTT |
| Anchor A_F | TCAGGAACATCGCTGACTCTCCGGACCAAGGGCTGAGGAGCAACAACAAA<br>C |
| Anchor A_R | TTCCGGTTGTATCCGGTAATGGTGACCAAGGCCTCAGCTCGTGTTCTG |
| Anchor T_F | GCCAAAGGCATCCATCTTCGTAACAGCTGAGGTCGTGTTCTTTG |
| Anchor T_R | CTTGTTGAAGATACGCAGGCCATTGCCTCAGCAGCAACAACAAAC |
| KLD nickRemoval F | TTTCAACGGCCTGCTCAA |
| KLD nickRemoval R | ATCAGCAAAACGGCGGTG |
| Dig-end competitor 1 | GGAGCAACAACAAACAGAAACAGACAAGAAACAAA |
| Dig-end competitor 2 | GCAAGAAACAAGAGAGAAACAGAACACGAGCTGA |
| Bio-end competitor 1 | GGTCGTGTTCTTTGGTTCTTTGTCTTGTTTCCTTG |
| Bio-end competitor 2 | TTTGTTCTTTCTTGCTTGTTTGTTGTTGCTGCTGA |
| 0.5 kb PCR F | TTATTCCAAGGGTAAAACGACGGCCAGTG<br>(underlined indicates the inserted StyI-HF site) |
| 0.5 kb PCR R | GGAAACAGCTATGACCATG |
| Dig-end competitor flip 1 | GCAGCAACAACAAACAAGCAAGAAAGAACAAACAA |

| | |
|---|---|
| Dig-end competitor flip 2 | GGAAACAAGACAAAGAACCAAAGAACACGACCTCA |
| 6.5 kb PCR F | Dig - GCCACCTGACGTCTAAGAAACCATTATTATCA |
| 6.5 kb PCR R | Biotin - GCTGGTCCTTTCCGGCAATCAGG |
| 12.7 kb PCR F | CTACCCGAGTGCGTGACAT |
| 12.7 kb PCR R | CCAGTCTCGTGAAGCGGTA |

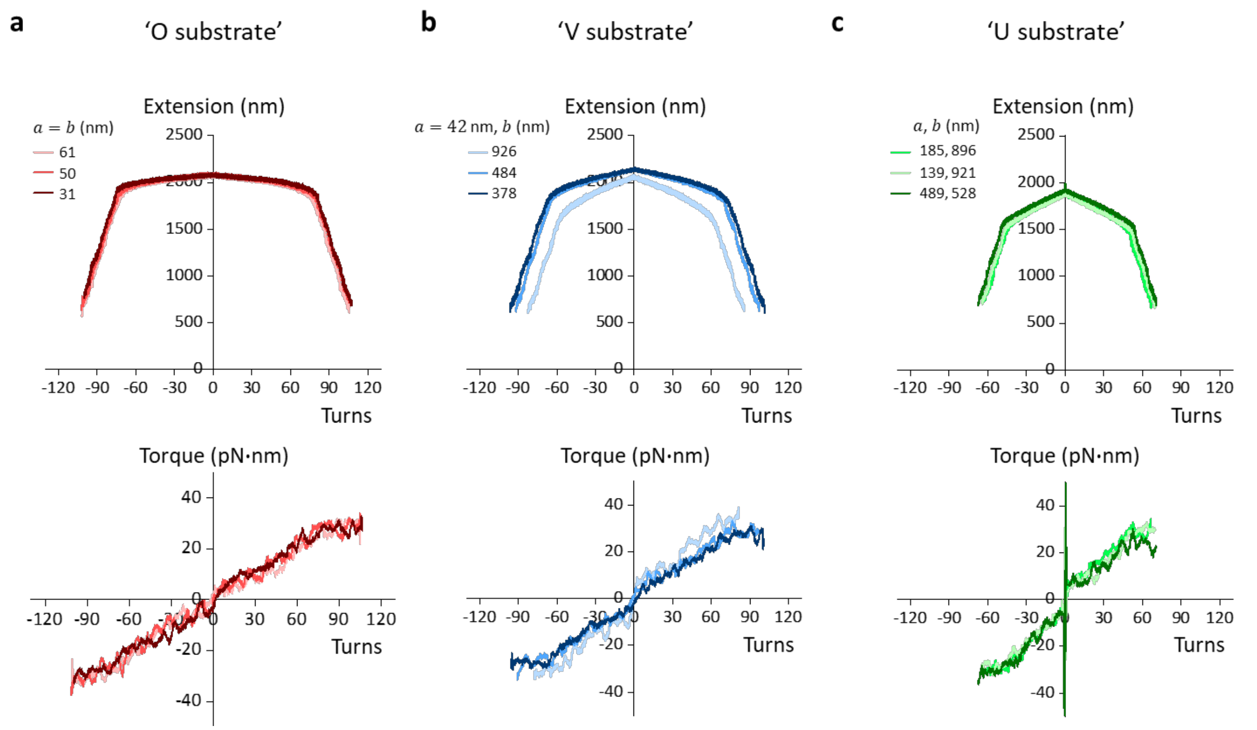


**Supplementary Figure 1. Individual traces of measured torsional responses during braiding the 'O substrate' (a), the 'V substrate' (b), and the 'U substrate' (c) at 2 pN.** For each braiding substrate, three example traces were presented to illustrate measurement and geometry uncertainties. The anchor separations of each trace are estimated by comparing the measured extension versus turns relation with those of simulations, as shown in Fig. 4b. Specifically, for the 'O substrate' traces, we assumed $a = b$, and the resulting values are close to the expected value of 42 nm from the simulations (Methods). For the 'V substrate' traces, we assumed $a = 42$ nm to obtain a more accurate measure of $b$. For the 'U substrate' traces, $a$ and $b$ are obtained directly. Source data are provided as a Source Data file.

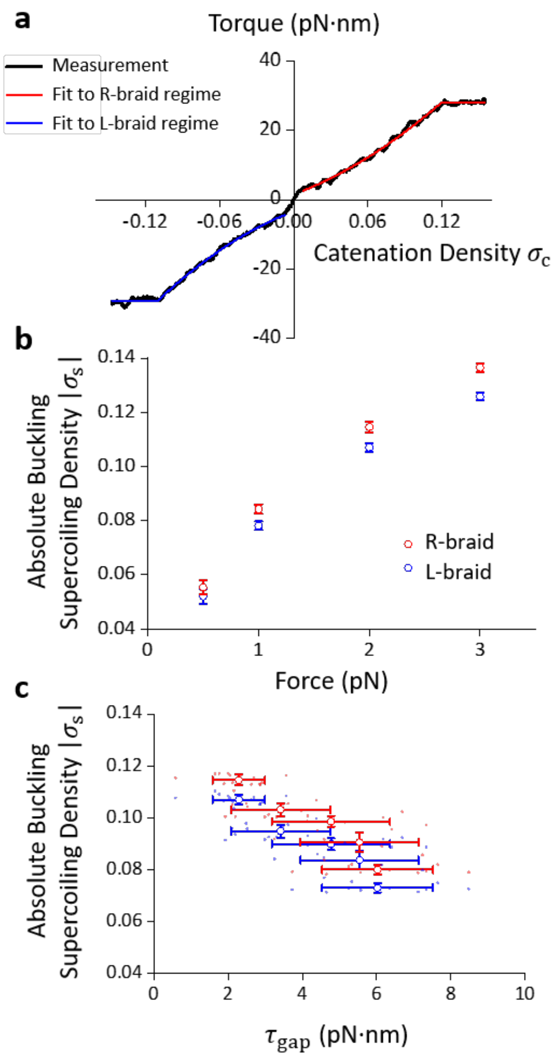


**Supplementary Figure 2. Chirality of DNA braiding.**

**a**. Determination of the twist-stiffening effect of R- and L-braided DNA. Shown in black is the measured torque versus supercoiling density from an average of *N* = 15 'O substrate' molecules held at 2 pN (same data from Fig. 2a). There is a small but detectable torque gap near the zero-catenation density. Outside this region, the measured torque profile can be well characterized by a 2nd-order polynomial function in the pre-buckling region and a constant in the post-buckling region.

For the R-braid, the fit (red) yields:

Braiding torque: $\tau = [(0.2 + 18.7\sigma_c + 101.3\sigma_c^2)\ \mathrm{nm}]k_B T\omega_0$, for $\sigma_c < 0.12$.

Braiding buckling torque: $\tau = (3.9\text{ nm})k_\mathrm{B}T\omega_0$ = 28.2 pN·nm, for $\sigma_\mathrm{c} \geq 0.12$,

where $\omega_0 = \frac{2\pi}{3.55\text{ nm}}$

For the L-braid, the fit (blue) yields:

Braiding torque: $\tau = [(-0.5 + 18.4\sigma_\mathrm{c} - 127.3\sigma_\mathrm{c}^2)\text{ nm}]k_\mathrm{B}T\omega_0$, for $\sigma_\mathrm{c} > -0.11$

Braiding buckling torque: $\tau = (-4.0\text{ nm})k_\mathrm{B}T\omega_0$ = -29.0 pN·nm, for $\sigma_\mathrm{c} \leq -0.11$

Source data are provided as a Source Data file.

**b**. Superhelical density at the buckling transition of the 'O substrate' at different forces (same data set as Fig. 2a). Each data point is shown as mean ± SD with *N* = 15 molecules. Source data are provided as a Source Data file.

**c**. Superhelical density at the buckling transition of braided DNA at 2 pN under different torque gaps (same data sets as Fig. 2a, Fig. 3a, and Fig. 3b). The data points from individual molecules are shown as a scatter plot (dots) and are further grouped and shown as mean ± SD. For the grouped data points *N* = 15, 12, 7, 5, and 11 molecules from small to large torque gap. Source data are provided as a Source Data file.

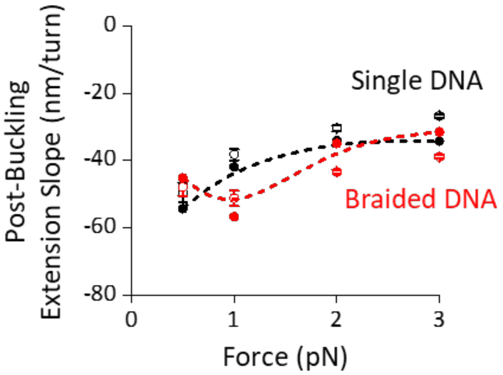


**Supplementary Figure 3. Post-buckling extension slope of braiding the 'O substrate' versus force.** Measurements (*N* = 15 individual molecules for braided DNA at all forces, and *N* = 20, 19, 18, 15 individual molecules at 0.5pN, 1pN, 2pN, and 3pN respectively) are shown as open dots with error bars as SDs and simulations as solid dots. The dashed lines are interpolations from the simulation results to guide the eye. Source data are provided as a Source Data file.

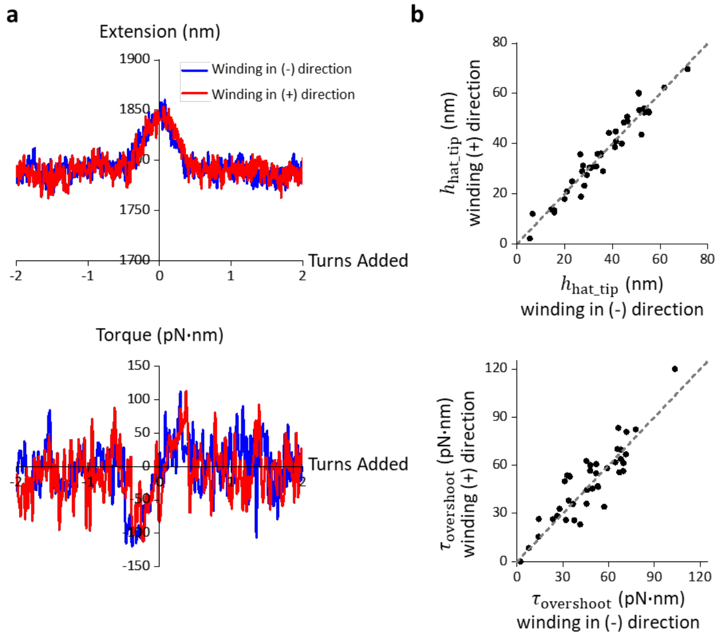


**Supplementary Figure 4. Reversibility of braiding with ±0.5 turns for the U substrate.**

**a.** Example trace of the extension and torque measurements within ±2 turns during (+) winding and (-) winding of the same tether. The extension-turns relations are similar for the two directions, and so are the torque-turns relations, indicating the winding to be near equilibrium. Source data are provided as a Source Data file.

**b.** Hat-tip and torque overshoot amplitude obtained from individual measurements of both winding directions of individual tethers (black dots). The measured $h_{\text{hat\_tip}}$ values from both directions of each tether are highly correlated, and so are the corresponding $\tau_{\text{overshoot}}$, further indicating winding to be near equilibrium. The grey dashed lines indicate the condition of full correlation. Source data are provided as a Source Data file.

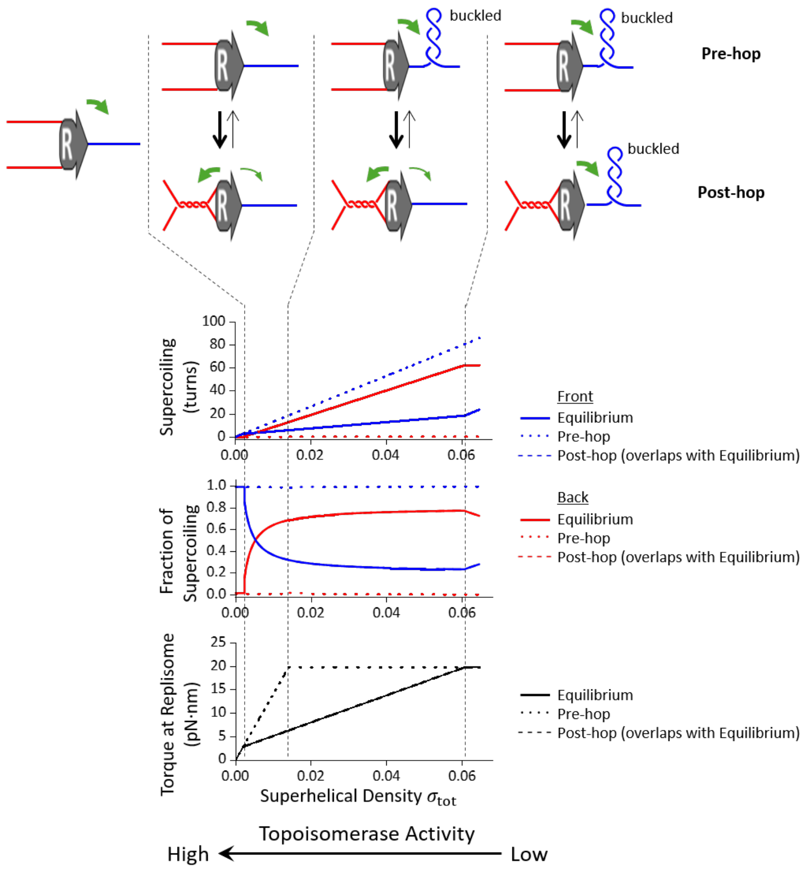


**Supplementary Figure 5. Supercoiling partitioning during DNA replication when both end separations are large.** Although this scenario is unlikely to occur during replication, we included this here as an extension to Fig. 5 for completeness. Here, we consider a braiding configuration ($a/l$ = $b/l \approx$ 0.14), similar to that of the 'U substrate', such that a torque overshoot is present. If the torque value rises above the torque gap but below the torque overshoot, the same torque value can be obtained at multiple catenation numbers, because the function of supercoiling versus torque is non-monotonic. We assume that different braiding configurations are in equilibrium and examine a thermodynamic ensemble of fixed total supercoiling across the forked DNA. Thus, the catenation number has a bimodal distribution peaking at a low catenation number before overcoming the torque overshoot (referred to as "pre-hop") and at a high catenation number after overcoming the torque overshoot (referred to as "post-hop"). We find the mean values for any state parameter (such as the catenation number in the braid and the torque of the forked system) using the partition function derived from the bimodal states.

For clarity, we also show the solution obtained by considering only the “pre-hop” state or only the “post-hop” state. Note that the “post-hop” and the equilibrium solutions nearly overlap, which is expected since the “post-hop” state has a lower energetic cost. The cartoons above the plots further clarify these states and how supercoiling partitions at each regime of the superhelical density. In the “pre-hop” state, supercoiling only partitions to the front of the replisome, whereas in the “post-hop” state, supercoiling partitions largely to the back of the replisome. The parameters used in this analysis are shown in Supplementary Table 3.

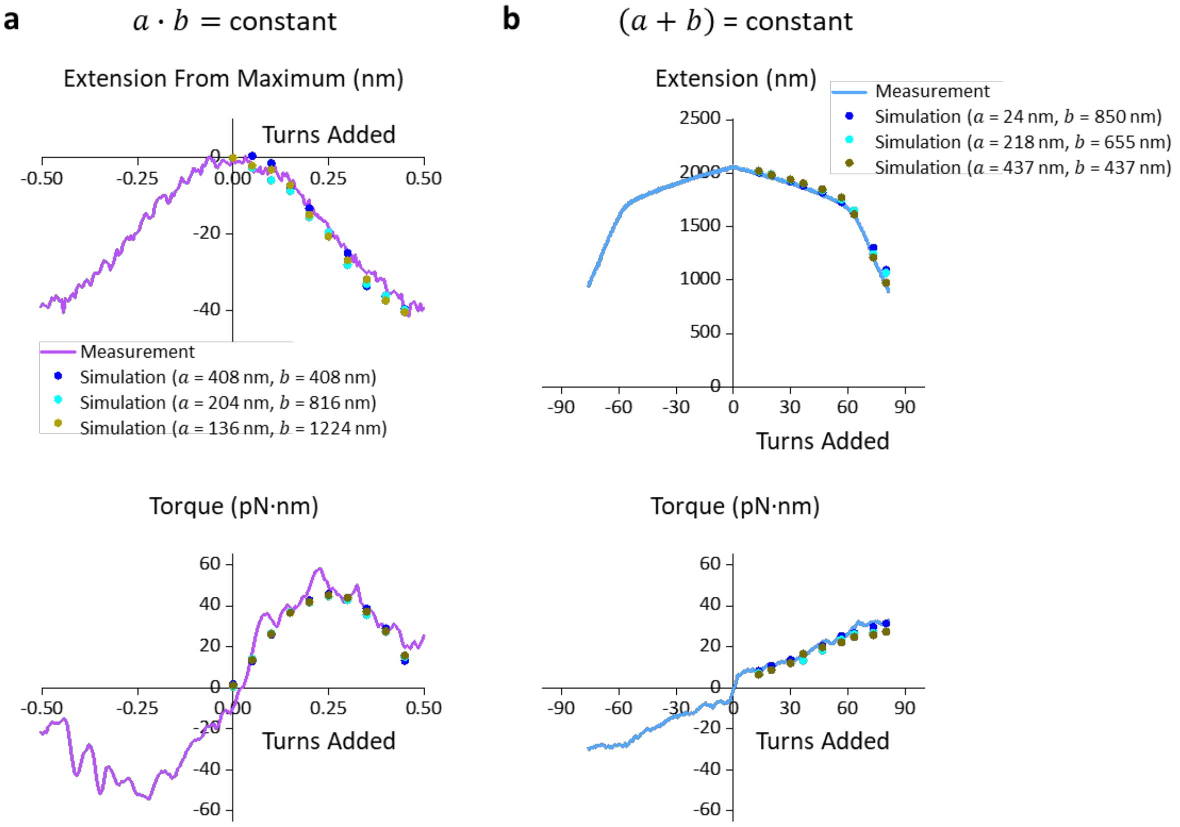


**Supplementary Figure 6. MC simulations of the extension-turns and torque-turns relations of braiding DNA with different combinations of end separations $a$ and $b$.**

**a**. MC simulations with different combinations of $a$ and $b$ while having the same $a \cdot b$ . Within ±0.5 turns, the extension-turns and torque-turns relations remain essentially the same for different combinations of $a$ and $b$. The measurements shown here are the same as those shown in the right plots of Fig. 3b. Source data are provided as a Source Data file.

**b**. MC simulations with different combinations of $a$ and $b$ while having the same $(a + b)$. Beyond ±0.5 turns, the extension-turns and torque-turns relations remain essentially the same different combinations of $a$ and $b$. The measurements shown here are the same as those shown in Fig. 3a. Source data are provided as a Source Data file.

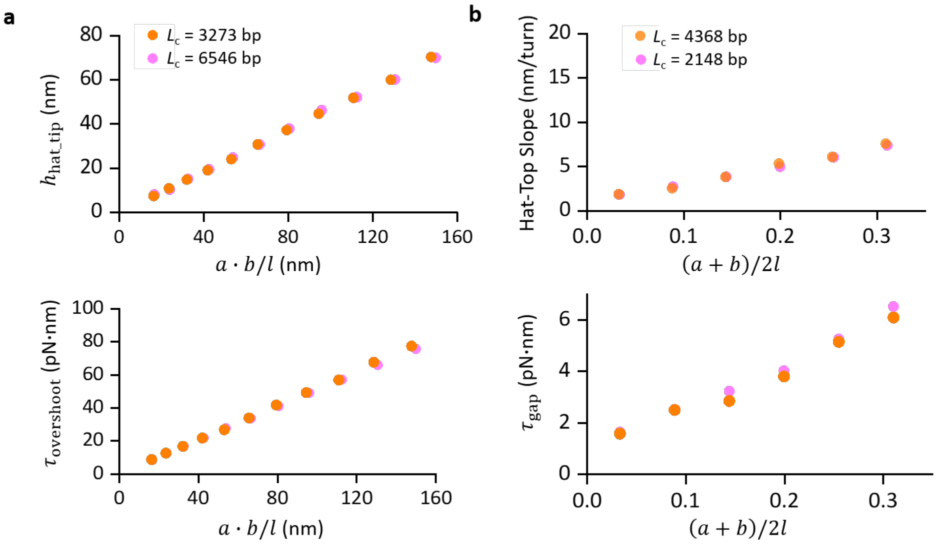


**Supplementary Figure 7. MC simulations of the torsional mechanics of braiding at 2 pN for different DNA lengths.**

**a**. Dependence of $h_{\mathrm{hat\_tip}}$ and $\tau_{\mathrm{overshoot}}$ on $a \cdot b/l$ for two different DNA contour lengths. Source data are provided as a Source Data file.

**b**. Dependence of hat-top slope magnitude and $\tau_{\mathrm{gap}}$ on $a \cdot b/l$ for two different DNA contour lengths. Source data are provided as a Source Data file.

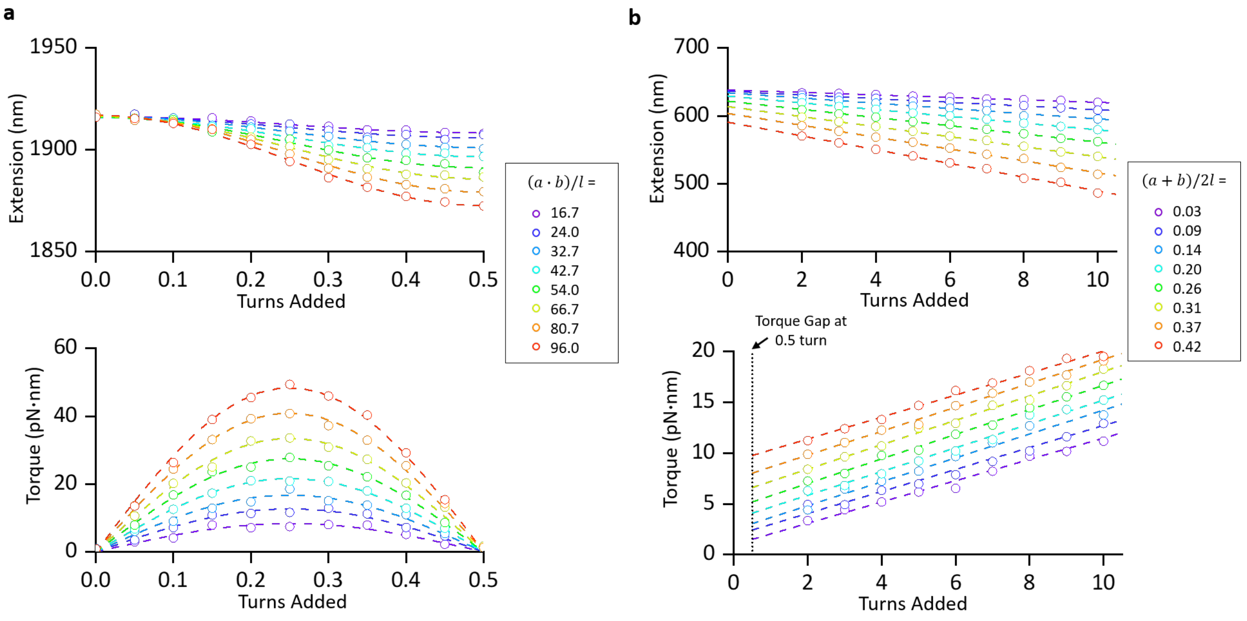


**Supplementary Figure 8. MC simulations at low catenation numbers for the U and V substrates with different geometries.**

**a**. The simulated extension and torque curves for *n* < 0.5 turns at different $(a \cdot b)/2l$ values (in nm). These plots are used for the hat tip and torque overshot analysis in Fig 4b. Dash lines are fits to guide the eyes. Source data are provided as a Source Data file.

**a**. The simulated extension and torque curves for *n* > 0.5 turns at different $(a + b)/2l$ values. These plots are used for the hat-top slope and torque gap analysis in Fig 4b. Dash lines are fits to guide the eyes. Source data are provided as a Source Data file.

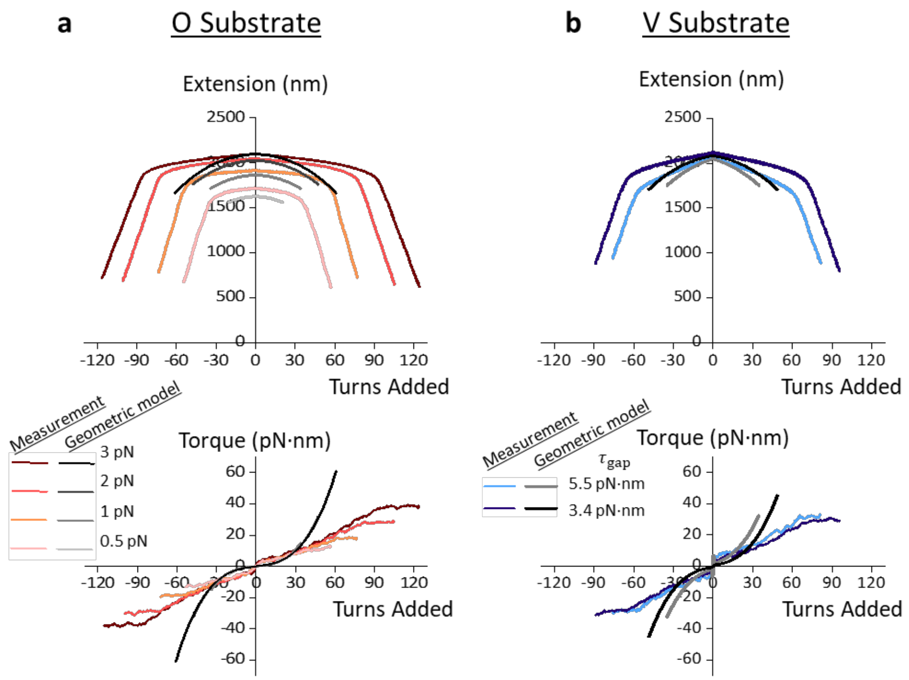


**Supplementary Figure 9. The geometric model does not accurately describe torsional properties once braiding has initiated.**

**a.** Comparison between the geometric model predictions and measurements under various stretching forces for the 'O substrate'. The separations at both ends $a = b$ = 47, 46, 42, and 41 nm at 0.5, 1.0, 2.0, and 3.0 pN, respectively, as determined by the MC simulations (Methods). Note that the predicted torque versus turns relations from the geometric model at the four forces nearly completely overlap. Source data are provided as a Source Data file.

**b.** Comparison between the geometric model predictions and measurements at 2 pN for the 'V substrate'. As determined by the MC simulations (Fig. 3a; Methods); $a$ = 42 nm at one end for both traces; $b =$ 850 nm for the light blue trace and $b =$ 421 nm for the dark blue trace, as determined from the MC simulations. Source data are provided as a Source Data file.

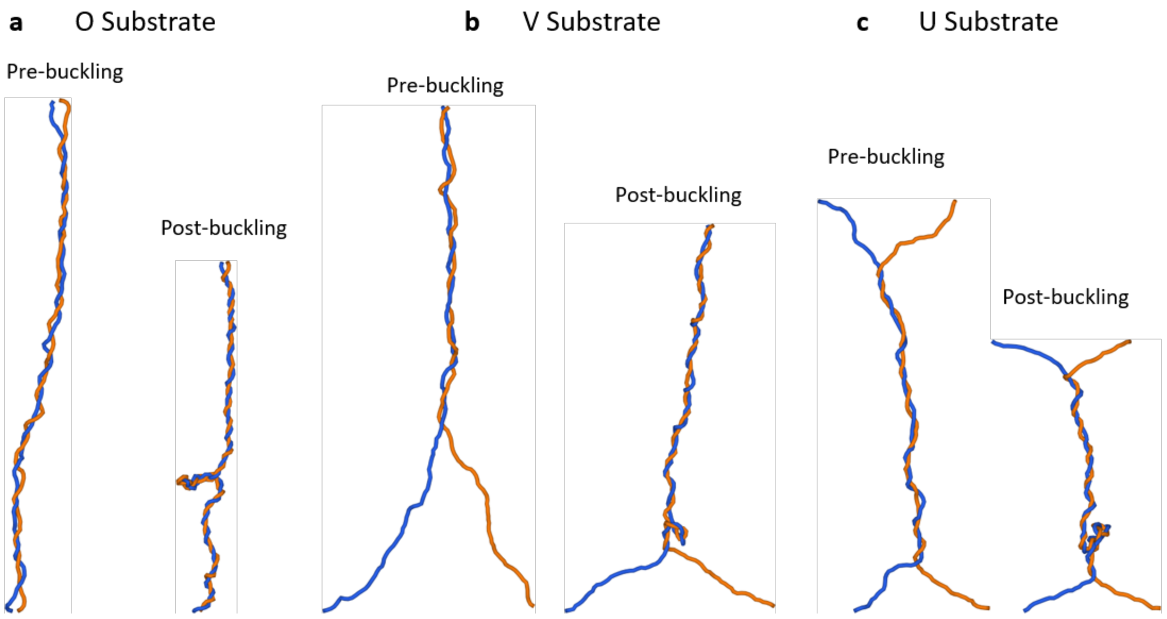


**Supplementary Figure 10. Configuration snapshots generated from MC simulations at 2 pN.**

**a**. O Substrate: $L_c$ = 2130 bp, $a$ = $b$ = 7.1 nm, 12 turns for the pre-buckled state, and 28 turns for the post-buckled state.

**b**. V Substrate: $L_c$ = 2184 bp, $a$ = 7.1 nm, $b$ = 255.0 nm, 6 turns for the pre-buckled state, and 19 turns for the post-buckled state.

**c**. U Substrate: $L_c$ = 1964 bp, $a$= $b$ = 162.8 nm, at 10 turns for the pre-buckled state, and 17 turns for the post-buckled state.

### Sequence of plasmid pMDW133

gatcctctagagtcgacctgcaggcatgcaagcttggcgtaatcatggtcatagctgtttcctgtgtgaaattgttatccgctcacaattccacacaacatacgagccggaagcataaagt
gtaaagcctggggtgcctaatgagtgagctaactcacattaattgcgttgcgctcactgcccgctttccagtcgggaaacctgtcgtgccagctgcattaatgaatcggccaacgcgcggg
gagaggcggtttgcgtattgggcgctcttccgcttcctcgctcactgactcgctgcgctcggtcgttcggctgcggcgagcggtatcagctcactcaaaggcggtaatacggttatccacag
aatcaggggataacgcaggaaagaacatgtgagcaaaaggccagcaaaaggccaggaaccgtaaaaaggccgcgttgctggcgtttttccataggctccgcccccctgacgagcatc
acaaaaatcgacgctcaagtcagaggtggcgaaacccgacaggactataaagataccaggcgtttccccctggaagctccctcgtgcgctctcctgttccgaccctgccgcttaccggat
acctgtccgcctttctcccttcgggaagcgtggcgctttctcatagctcacgctgtaggtatctcagttcggtgtaggtcgttcgctccaagctgggctgtgtgcacgaaccccccgttcagcc
cgaccgctgcgccttatccggtaactatcgtcttgagtccaacccggtaagacacgacttatcgccactggcagcagccactggtaacaggattagcagagcgaggtatgtaggcggtgc
tacagagttcttgaagtggtggcctaactacggctacactagaagaacagtatttggtatctgcgctctgctgaagccagttaccttcggaaaaagagttggtagctcttgatccggcaaa
caaaccaccgctggtagcggtggtttttttgtttgcaagcagcagattacgcgcagaaaaaaaggatctcaagaagatcctttgatcttttctacggggtctgacgctcagtggaacgaaa
actcacgttaagggattttggtcatgagattatcaaaaaggatcttcacctagatccttttaaattaaaaatgaagttttaaatcaatctaaagtatatatgagtaaacttggtctgacagtt
accaatgcttaatcagtgaggcacctatctcagcgatctgtctatttcgttcatccatagttgcctgactccccgtcgtgtagataactacgatacgggagggcttaccatctggccccagtg
ctgcaatgataccgcgagacccacgctcaccggctccagatttatcagcaataaaccagccagccggaagggccgagcgcagaagtggtcctgcaactttatccgcctccatccagtcta
ttaattgttgccgggaagctagagtaagtagttcgccagttaatagtttgcgcaacgttgttgccattgctacaggcatcgtggtgtcacgctcgtcgtttggtatggcttcattcagctccgg
ttcccaacgatcaaggcgagttacatgatcccccatgttgtgcaaaaaagcggttagctccttcggtcctccgatcgttgtcagaagtaagttggccgcagtgttatcactcatggttatgg
cagcactgcataattctcttactgtcatgccatccgtaagatgcttttctgtgactggtgagtactcaaccaagtcattctgagaatagtgtatgcggcgaccgagttgctcttgcccggcgt
caatacgggataataccgcgccacatagcagaactttaaaagtgctcatcattggaaaacgttcttcggggcgaaaactctcaaggatcttaccgctgttgagatccagttcgatgtaacc
cactcgtgcacccaactgatcttcagcatcttttactttcaccagcgtttctgggtgagcaaaaacaggaaggcaaaatgccgcaaaaaagggaataagggcgacacggaaatgttgaa
tactcatactcttcctttttcaatattattgaagcatttatcagggttattgtctcatgagcggatacatatttgaatgtatttagaaaaataaacaaataggggttccgcgcacatttccccg
aaaagtgccacctgacgtctaagaaaccattattatcatgacattaacctataaaaataggcgtatcacgaggccctttcgtctcgcgcgtttcggtgatgacggtgaaaacctctgacac
atgcagctcccggagacggtcacagcttgtctgtaagcggatgccgggagcagacaagcccgtcagggcgcgtcagcgggtgttggcgggtgtcggggctggcttaactatgcggcatc
agagcagattgtactgagagtgcaccatatgcggtgtgaaataccgcacagatgcgtaaggagaaaataccgcatcaggcgccattcgccattcaggctgcgcaactgttgggaaggg
cgatcggtgcgggcctcttcgctattacgccagctggcgaaagggggatgtgctgcaaggcgattaagttgggtaacgccagggttttcccagtcacgacgttgtaaaacgacggccagt
gaattcgagctcggtacCGGAGGATGGCAGCGTGATTTCACGGTCGAGCGTCAGCGTCCGGGTCTGGCTGTTCACCGCCAGCACACGACCACCGG
TGCTGATACCGGCATAGTCATCATCGCAGATTTCAATAACATCGCCCGGTACATGGCGAAGCCCTTCTGCGCCGACGCTGAAATCCACGGTCT
GCGTTTCCAGCAGTTCTGTTTTAATCAGCCACAGCCCGGCGCGGTGTGCCTGCCCCCGGCTGGTACAGCCAAAGGCATCCATCTTCGTAACAT
TACGACCGTAACGGGCAATGGCCTGCGTATCTTCAACAAGCTCTGTCGCCGTCTCCCAGCCGTTGTTCGGGTCAATCCAGTTCACCTCAACGG
CATTATGGCGGTCCTTCAGGGCGCTGAAGCTGTAGCGGAACGGCGCGCCATCATCCGGCATCACCACATTACTGCGGTTATAGGTCCACGTCT
TATCCGACGGTCGGTCCTGCACGAACGTCAGCGTCTGCCCGTTCCATACCGGCATACAGCGCATCGCCGAGCAGAAATCGCTGAGCACATCC
CACGCCTTACGCTGTGTGGTCAGGTACGCATTACAGGTGATGCGCGGCTCCGTGCCGCCAAAGCCGTCCGGCACTGACTGGTCGCAGTACTG
GCCGATGACATACAGCGCCCATTTATCCACATCCGCCGCACCAAGACGTTTCCCCATGCCGTAGCGCGGATGGGTCAGCATATCCCACAGACA
CCAGGCCATGTTGTTGCTGTATGCCGGTTTAAACGTTCCGTCCCAGATACCGCTGTATTGCCGCGTCTGCGGGTTATAGTTCGACGGCACCTG
CAGAATACGCCCGCGCAGATGATAATTACGGCTCACCTGCTGGCTGCCGAACTGCTCCGAGTCCACCTGCACGCCGACCAGTGCCGTGTTCG
GGTAGCACTGTTTCACATCGATGATTTCAGTGTATGACGACCAGAGCGTTTTGTTCTGCAGCTGGTCTGTGGTGCTGTCCGGCGTCATCCTGC
GCATCCGGATATTAAACGGGCGCGGCGGCAGGTTACCCATCACCACCGAGGCCAGATACTGCGAGGTGGTTTTGCCCTTAATGGTGATGTCT
TTTTCCGTCACCCAGCCACCGTTACGTTGTATCTGAACCAGCAGGCGGACTTCCGACGGATTCCTGTCACCCTTTGAGGTGGTTTCCACCAGTG
CCTGTACACCGAAGGTAAAGCGCAGACGGTCGATGTTTGCAGACGTAATGGTGCGGGTGATCGGCGTGTCATATTTCACTTCCGTACCCAGC
ACCGTCTCGGAGCCGGAGGATTCAAATCCCTCCGGCGGAGTCTGCTCCTGCTCACCAGCCCGGAACACCACCGTGACACCGGATATGTTGGT
ATTCCCCTCAGTGTCCAGCACCGGCGTACTGTTCAGCAGCACGCTTTTTAAGCCATCCACCGGACCTTCAATCGGCCCTTCGCTGATGGCATCG
ATCACACTCAGCAACTGCGTGGACTTCAGGTTGTCCTTCGCTTCGCGCGGGGTATGCCCCTTACTGCTTCCTTTACCCATTCCTCACGCTCCATA
AATGACAAAACCGCCCGCAGGCGGTTTCACATAAAACATTTTGCATCAGCGACCAATCACCACAACCTGACCACCGTCCCCTTCGTCTGCCGT
GCTGATCTCCTGAGAAACCACGCGTGACCCCACGCGCATTTCCCCGTACAGAACAGGCAGAACATTGCCCTGGGCAACCATGTTATCCAGTGA
GGAGAAATAGGTGTTCTGCTTACCGTTATCCGTTGTCTGTATACGGGGAGTTCTGGCTTTCGGTGCCAGCATCTGCGCCACACCACCGAGCAC
CATACTGGCACCGAGAGAAAACAGGATGCCGGTCATACCACCGGCCCCAATGGCTGCCCCCCATGCTGCAAGGGTGGCTCCGGCGGTAAAG
AATGATCCGGCAATGGCGGCAGCCCCCAGGACAATCTGGAATACGCCACCTGACTTGGCCCCGGCGACTCTGGGAACAATATGAATTACAGC
GCCATCAGGCAGAGTCTCATGTAACTGCGCCGTTAACCCGGACGTGCTGACGTCCCGCCCGGCAATCCGTACCTGATACCAGCCGTCGCTCAG
TTTCTGACGAAACGCCGGGAGCTGTGTGGCCAGTGCCCGGATGGCTTCAGCCCCCGTTTTCACACGAAGGTCGATGCGGCGACCAAATCGTT
GTAAATCCCCGTAAAGGCAGATGCGCGCCATGCCCGGTGACGCCAGAGGGAGTGTGTGCGTCGCTGCCATTTGTCGGTGTACCTCTCTCGTT
TGCTCAGTTGTTCAGGAATATGGTGCAGCAGCTCGCCGTCGCCGCAGTAAATTGCGGCGTGATTCGGCACTGATGAACCAAAACAGCACAGC
AGCACATCGCCCGGCTGTGCCGCTGACAACGGCACCTGATACAGCCCCGTCGCCTCCAGATTATCCAGATAGAGATTCTGGCCGTTACGCCAC
CAGTCATCCTCACGATGAAAGTCCGGCATCTCAATCCCCGCCAGATGATAAGCATCCCGGAACAGTGTGTAACAGTCCGTCACACCGTGCTCA
AAGCGCCGCCCGGTGAGATGCGGCACACAGCGGAACTTATGAATCGTCCCCCGGCAGACCAGCCACCACGGCAAATCACTCTGCACCTGCAG
CCGCCGGTCGGCCTCACTCAGCCAGGGCAGACCACCGGGGTGGCTGTGGACCAGCGCCACAATCTCACCCTGCATTTCTGCCTGCAGCCAGT
CTTCCGGCGACATACGGAAATAGCCTCCGGCTCACCGGAGATATTCACGCAGGGGAAATATCTTTCCCCCTCCGGCGTGCTTACCACGAAGCC
GCACGACTCCGCTGGCGCACATCGCCGGGCGTGCGCCAGAATCGCTGATTCTGTCTGTGTCATGGGATTTACTGCGAAAGTTTGTTAATGGA

AAGGAAGCCGCCAAAGTTGCCGACGTTATTGCGGAACTTACAACCGCTCAGGCATTTGCTGCATTTATCCTTCGTGATATCGGACGTTGGCTG
GTCATATTCATCCGCGACAGCCGGACCGCTATAACCGCACTCGTCACCGCGATAGGTCCAGGTGCAGGTGTTGGCCAGCATGATACGTCCCG
GAAAAACAGCGCCATCCGTTTCCGTCGGCGTGGACAGTACAAAGGAGGCACTCACCGCGCTCAGTTCGCTGCACTGCTCAATGCGCCAGCGG
CTGATCACCTCCTGCTCCGGATCGGCGTAACTGTTTCCGTTGACGAAGTTCACCGCATCCAGAAAACGGGCGTAAACCTTACGCCGGACCACC
GTTCCGCCGACCAGACTCTGCATATCTTCCGCCATCCCGGTGACCATACCGTACAGGTTAGAAACCGTCAGCGTGGGGCGCGTACTGGTGCCT
TTGCCATTCAGTTCAAAACCGCTCCCCTGAATGGGATACGGCTGATACTGTCGCCCCTGCCAGGTGACCGGCTCACCTTTTTCGTTCTGCTCAT
TACAGAAAAAATAACGTTCTCCACCGACCTCTGTCAGGTCGATTTCCCAGAGCACCACGCTGGCCGACTGCTCCGCACGGGTGCATTCATTCA
GTGTTTCCTGCCGGATATCCTGCATCAGTTCACCACCTGTTCAAACTCTGCGCTGAACTCAACACGCAGCATACTGACCCGCGACGACCATTTT
GCGCAGGTCACCTTTATCTGCCGCCACTCATAAGGCGGCGTCCACAGAAAGGATTTCCAGCCCCCGTGCTCTTCCAGAAACGACTCCAGTACC
GTGGCCTCCTCACGGGGGACAGAAAGCGTCACGCTGTACGTTTTCAGGTTGGCATTCAGCCCGGCAGGCGCTCGCTGAGAATAGCCATCACC
AAAGCGCACCTTTCTTACAGAAGGGACCGAAGCCACATCCATACCGGGTTTCACTTTCCAGCGGAAGGTCTTCATCGTCCACCTCCGGAGAAC
AGGCCACCATCACGCATCTGTGTCTGAATTTCATCACGGGCACCCTTGCGGGCCATGTCATACACCGCCTTCAGAGCAGCCGGACCTATCTGC
CCGTTCGTGCCGTCGTTGTTAATCACCACATGGTTATTCTGCTCAAACGTCCCGGACGCCTGCGACCGGCTGTCTGCCATGCTGCCCGGTGTAC
CGACATAACCGCCGGTGGCATAGCCGCGCATCAGCCGGTAAAGATTCCCCACGCCAATCCGGCTGGTTGCCTCCTTCGTGAAGACAAACTCA
CCACGGTGAACAATCCCCGCTGGCTCATATTTGCCGCCGGTTCCCGTAAATCCTCCGGTTGCAAAATGGAATTTCGCCGCAGCGGCCTGAATG
GCTGTACCGCCTGACGCGGATGCGCCGCCACCAACAGCCCCGCCAATGGCGCTGCCGATACTCCCGACAATCCCCACCATTGCCTGCTTAAGC
AGAATTTCTGTCATCATGGACAGCACGGAACGGGTGAAGCTGCGCCAGTTCTGCTCACTGCCGGTCAGCATCGCCGCCATATTCTGTGCAATA
CCATCAAAGGTCTGCGTGGCTGCACTTTTTACCTGCGACATACTGTCCGTGGCGCTCTCTTCCCACTCACTCCAGCCGGACTTCAGGCCTGCCA
TCCAGTTCCCGCGAAGCTGGTCTTCAGCCGCCCAGGTCTTTTTCTGCTCTGACATGACGTTATTCAGCGCCAGCGGATTATCGCCATACTGTTC
CTTCAGGCGCTGTTCCGTGGCTTCCCGTTCTGCCTGCCGGTCAGTCAGCCCCCGGCTTTTCGCATCAATGGCGGCCCGTTTTGCCCGTTGCTGC
TGTGCGAATTTATCCGCCTGCTGCGCCAGCGCGTTCAGGCGCTCCTGATACGTAACCTTGTCGCCAAGTGCAGCCAGCTGGCGTTTGTACTCC
AGCGTCTCATCTTTATGCGCCAGCAGGGATTTCTCCTGTGCAGACAGCTGGCGACGTTGCGCCGCCTCCTCCAGTACCGCGAACTGACTCTCC
GCCTTCCACAAATCCCGGCGCTGCTGGCTGATTTTCTCATTTGCTCCGGCATGCTTCTCCAGCGTCCGGAGTTCTGCCTGAAGCGTCAGCAGG
GCAGCATGAGCACTGTCTTCCTGACGATCGCCCGCAGACACCTTCACGCTGGACTGTTTCGGCTTTTTCAGCGTCGCTTCATAATCCTTTTTCGC
CGCCGCCATCAGCGTGTTGTAATCCGCCTGCAGGATTTTCCCGTCTTTCAGTGCCTTGTTCAGTTCTTCCTGACGGGCGGTATATTTCTCCAGC
GGCGTCTGCAGCCGTTCGTAAGCCTTCTGCGCCTCTTCGGTATATTTCAGCCGTGACGCTTCGGTATCGCTCTGCTGCTGCGCATTTTTGTCCT
GTTGAGTCTGCTGCTCAGCCTTCTTTCGGGCGGCTTCAAGCGCAAGACGGGCCTTTTCACGATCATCCCAGTAACGCGCCCGCGCTTCATCGTT
AACAAAATAATCATCCTTGCGCAGATTCCAGATGTCGTCTGCTTTCTTATACGCAGCCTCTGCCTTAATCAGCATCTCCTGCGCGGTATCAGGA
CGACCAATATCCAGCACCGCATCCCACATGGATTTGAATGCCCGCGCAGTCCTGTCTGCCCAGGTCTCCAGCGTGCCCATGTTCTCTTTCAGGC
GGCGGGTCTGGTCATCAAACCCTTTCGTTGCGGCCTCGTTCGCCGCCTGCAATGCCCCGGCTTCATCGCCGGAACGCTGCAACTGAGCAACAT
ACGCAATCTGCTCCGCCGACACGTTATGGAACTGGCGAGCCATCGCCGTCAGCCCCGACGTCGGGTCTGTGGTCAGCTTCCCGAAGGCTTCA
GCGACCTTGTCCACCTCCACGCCGGATGCAGAGGAGAAACGCGCCACACTCTGGCTGATGGACGCAATCTGAGCCTCACCGCTTACCCCCGC
CTTAACCAGTGCGCTGAGTGACTCGCTGGTCTGGTTAAACGTCAGCCCTGCCGCCTGCCCGGCTCTGGACAGGACCAGCATACGATCTGCCGT
CAGTCCCGCCTGATTGCCGGAAAGGACCAGCGTTTTGTTGAAATCGGACAGGGTTGAGTTGCCCTGATACCAGGCATACGCCAGCGCACCGG
TCGCCACCGCCAGCGAGGTGGCCCCCACCATCGGCAGGGTGATCGCACCGGCAAGCCCCCTGAACATGGGGATCATCCCGCCGAAGGAGTC
CTTCACCTGCCCCCCCTGTTGCAGCAGGATCAGCCACGGACTTTGCCCGCCTGCAAGCTGCGTGGCCACGTCGGTGAACTGTGCAGGCAGCAT
ACGCATGGCGGCTTTATACTGCCCGACGGAAATCCCCGCTTTCTGTGCAGCCAGCGCCTGTCGGCTCAGCGACTGTTCAACGACTGCCGCTGT
TTTTTTCGCATCACTTTCCGTACCAGAAAAATGACGCCTGACTCTGGCCATCTGCTCGTCAAATCTGGCCGCATCCAGACTCAAATCAACGACC
AGATCGCCTACCGGTTCAGCCATACCGGACTCCTCCTGCGATCCCTTCTGATACTGTCATCAGCATTACGTCATCCTCCGTCATGTCCGCCACAT
CCGGGGAAGCGGGGATAACTTCATTCCCGTCCGGGCCAAAGCGGACACCTCCGGCAAGCCCTGCCGCTTTCTGCATCAGCACATCATCTTCA
GGCTCTTCGTCAGCCTCGCGCCGGTTCAGCAGACTGAAATCCAGCGGATGCATATCCGGATCGCTGAAAAACAGGCTGAGCACGGTGTACGT
CAGCCCGGAAAAGTGCATATCCAGCAGAACATCATGAAAATAATGGGTACTGTAAAAGCGGTGCCAGTCGGCATACTCCGTGGATGACATCC
CGGCAAGCATGGCACGCCAGTCGGGTCGCCCCATCTCACGCGCCAGTTTCAGGGCAAAACTCAGCTCACCGTCGAACACTTTCCCGCAGAAA
CAGGCTCTGCGGGCCCGGCGTCCTCTGTCTGTTCAGGGGCATTATTCACCACAAACTCATACATACCAGACAGCCGGTACACCACGTTTTCAG
CATGAGAAATTGCCTCCGTGGGCCAGGTGGTAAGCACTTCCTGCTCAATCTGTTTAACGGCTTCATTCATGGACGGCATCTGCGTCTTCTGCG
GATGGTTATGCCACAGGGACATCGCCACCAGAAACGCGCCGGTTCTGATGGCGTCTTCCACAGTAAACTTCCGGTTGCTGTCTGACTCCGCCT
GTTCTGCCTGCCGTTTCATCAGGGCGAGATGCTCAATGCGCTGCAGGGCTGACAGTTCAGAAAGCGTGACGGTCACACCGTTATGTTCAAAT
GATTCGGTTTTCAGGAACATCGCTGACTCTCCGGATTAACTGGCGGTGACGGTAATTTCTGCAACCGCAGCAAACTCACCATTACCGGATACA
ACCGGAATGTTGACCTTGCCTGCAGCAACGCCGTTCACGGTGATGGTCATACCACTGACCGACACGGTGGCTTTTGTTTTATCCGCAGACACC
GCACGAAAGCTCTTGTCGGTTACGCCCTCCGGCTGGAAGGCCACGGTCAGCGTGGTGCTCTGCCCTTTCACCACCGAGGTGCTGGCAGGCGT
CACGGTCATGCCGGTTGCCGCTGTTACCGTGCTGCGATCTTCTGCCATCGACGGACGTCCCACATTGGTGACTTTCACCGTGCGGGTGATCAC
TTCCTTCGCCGTCACCGCCTTACCGATACTGCTGACCCAGCCACGGAACACATCGACCGTGCCGTTCGGGAAGCGGATTTTATAGGCACGGGT
ATCGCCTTCATTAAACCACGCCAGCAGCGCCTGCTGCCCCTGCTCTCCGGGCATCCACGCCAGCGTGAAGCTGGTATCTCCGGCAGATTTCTG
CCCCTGCCCGGTCGCAGTCCAGTCTGCATCTTCATCATCGAGATAGCTGTCGTCATAGGACTCAGCGGTCAGTTCGCCGGGCGTCAGGTCTTT
AACTTTTGCCAGACGCGACCAGTCAACGTCTGAAAGCGGATTCGCGTAAGGGTCACCGCTCCCCTTATAAACCCACAGGGTGGTCCCGGCAC
CTTTCACCGGCATTGTAGGATTTGGTACAGGCATAGCGTCCTCACATTTCATAGGTAATGACATAAGTCAGATCGGCTGAACTCCACAAGCCC
GCATCATCGTCGCGCCGGTAGTCATAGCCGCTGGCCACCATACTGGTGATCAAATCTGACAGTGCCGGGATATCGCTCATCACCGGATAAATC
CGGGACTCCATCCACGCATCCAGCTCTGAATCCGGCACCTGAGCAGGCAGGAAAACTTCGATATGCAGCTCCGCCTGCCAGGTATCGCTGTCC

AGCTCTTCGCCCGTGTATTCAGCGCCGGTGAGATAAACGGCAACTGCCGGAAAATCCGCCTCATCAAAAACAGCGGGGCGACCATCAAAAAA
CGTCGCCCCGGTGTCATGCTTCTCCAGTGCATCCAGTACGGCTGCACGGAGTTCAGTATGTTTCATCGCTTTATTACCATCCTCAGTTGATGCT
GCAGCGCATAGCCCAGCTCTTTCGGAAGACGTTCACGCCGTATCCGCTCAATATTTTGTTTAAACGCCGTGGTCAGCGGCACCGCCATCGGGA
TTTTCACCACATCAATGGGGTAACGGTTTTTCCCAGCCACACGCTGCATGACATGCCACCGGCCATTTTTCAGTTGCTGAATAAACGCGCCGG
GAATACGACGGTTACCCACCACAAGCACGCTGCCGCCACCTTTCAGGGATGAACGCTGCCCCTTTTTACGACGCCTGCGGCGCGAAAGGACA
ACCCGCGCATTACCCAGCTTGATTACGGGCAAATCCCCCCGGTTAACTTTGATTCTGGCCTGCGGATTTTTGACCGTGGCCCTTTTCAGCCTGG
CCCTTTCCTTTACCAGTTTCCGGCGTACCTTTGTCTCACGGGCAACCTGTGACGCCGACTGCGATATCGCGGATGAAGCAACGCGGTTAATGG
CCATTGCGGCGGCACCAGGCACCGCCGTTTTGCTGATACGGCTGAGGTTTTCAACGGCCTGCTCAAGACCTTTTATGGCCATACATCCCCCTTT
CAGCGGCGACGGTTAACGGCAGGCGGTACGCCCCGTCCAAGCCAGAGATGACAACTTCCGCCATCATCCGGCGAAACCCGATCTACCCAGAA
ATTTTCCTCACCGATGGTCAGCGTGTCTCCACGCCGCAGCTGCCGCACCTCATCAGTCCGGACAAACAGGGACGGGCTGGAGCCTTCAACGC
GCACGCCCTGTCCGGCATAGCTGATATTTTCAGGGTCATCAAAAACACCACGTATCACCGCACCTGACTGCTCACCGGATGTAATGGTGGCTG
ACGTTCCCATGTACCCGCGTATCGTTTCATCGGCGCGGGCAATGGCAGCATCGAACAGGTTATCGAAATCAGCCACAGCGCCTCCCGTTATTG
CATTCTGGCCAGGCCGCGCTCTGTCATTTCGGCTGCCACACCGGCAGAGACACGAAACGCCGTTCCCGGCAGCACAAATGCCACAGGTTCAT
CCCGCGTGGCGTGAAGTGCATCAGTATGCAGCTTCACCAGTGCCACGACCGTGACCAGTTCAGACGTATCCAGAATCACGGTATCCGGCTGC
GCTGATCCCACCTCATTTTCATGTCCGGTCAGCACATTTTCCCGGCTGAGAGGGGTGTCCTGACCGGCAGTTTCATCCGTGTCATCAAGCTCCT
CTTTCAGCTCTGCCACACGGAGCGCCAGTTCTTCTTTCGTCCCCGTCAGGCTGACATCACGGTTCAGTTGTTCACCCAGCGAGCGGAGACGGG
CAATCAGTTCATCTTTCGTCATGGACTCCTCCACAGAGAAACAATGGCCCCGAAGGGCCATGATTACGCCAGTTGTACGGACACGAACTCATC
AGGGTCAGCCAGCAGCATCAGCGGTGCTGACTGAATCATGGTGAACTCACGCGCCGGATCGCCGGTGGTCACCCAGTTTTTCGGGTAACGG
GCAGAGGCGTTAATGCCTTCGCGCTGTGCGTCCGCATCCTGAATGCAGCCATAGGTGCGCAGACCGCGTGCCTGAGTGTTCCCCAGCACCAT
CGTGTTGTCCGGCAGGAAGTTCTTTTTGACGCCGTTTTCCACGTACTGTCCGGAATACACGACGATGGCCACATCGCCATACATCCCCTTATAG
GACACCGCTTTGCCCAGGTCTTTCACCGCTGTCTCCAGCTCGGAATTAGAGCCACGACGGGTATCCAGCTTCTCCTTGACGGCTTTGAAGGAA
CGGAACAGCGCCCAGCCTTTCGGATCGAACACGATGATATTCACCACACCGCTGGCGTTCAGCGCGTAGGCTTCGATATCGTCGGTCGGGTC
ATACGTGGACTTGTCACGCTTGCTCCACTCCGTGCCGCCGGACTGCGTGATGTTATTCTCCTCACTGCGGCCCATATCCACCTCAACCGGATCG
AAGGCTTCACCGGTCATGGTGTATTTGCCCTTAAGCACGGCAGAAACTGCCTGCATCTCTTCGACCTGAGCAATGGCCAGCTCTTCGTCACGC
ATGTTCTGCATGATGATGCGACGGCGGCGGTAAGCCGGGTCCGCCAGATTCTGCGGATCTTCATCCGGCAGGCGACGCAGGGTCATCTGCG
GATTCACTTCATGCTTCGGCTTGACATATCCCGGCGTAAATTCAGAGGTGGAGCCGCCACGGGAACGGATAACCTCACCGGAAACAATCGGC
GAAACGTACAGCGCCATGTTTACCAGTCCCGGAATTTGTGAGAGATAGACTTTCTCCGTGGTGAAGGGATAGCTCTCACGGAAAAAGAGACG
CAGAAACAGCGGATCAAACTTAAATTTCTGCTCATTTGCCGCCAGCAGTTGGGCGGTTGTGTACATCGACATAAAAAAATCCCGTAAAAAAAG
CCGCACAGGCGGCCTTTAGTGATGAAGGGTAAAGTTAAACGATGCTGATTGCCGTTCCGGCAAACGCGGTCCGTTTTTTCGTCTCGTCGCTGG
CAGCCTCCGGCCAGAGCACATCCTCATAACGGAACGTGCCGGACTTGTAGAACGTCAGCGTGGTGCTGGTCTGGTCAGCAGCAACCGCAAG
AATGCCAACGGCAGCACCGTCGGTGGTGCCATCCCACGCAACCAGCTTACGGCTGGAGGTGTCCAGCATCAGCGGGGTCATTGCAGGCGCTT
TCGCACTCAATCCGCCGGGCGCGGTTGCGGTATGAGCCGGGTCACTGTTGCCCTGCGGCTGGTAATGGGTAAAGGTTTCTTTGCTCGTCATA
AACATCCCTTACACTGGTGTGTTCAGCAAATCGTTAACGGCATCAGATGCCGGGTTACCTGCAGCCAGCGGTGCCGGTGCCCCCTGCATCAGA
CGATCCAGCGCAGTGTCACTGCGCGCCTGTGCACTCTGTGGTGCTGCGGCCAGAATGCGGCGGGCCGTTTTCACGGTCATACCGGGGGTTTC
TGCCAGCACGCGTGCCTGTTCTTCGCGTCCGTGAGCCTCCTCACAGTTGAG

### Sequence of plasmid pMDW168

AAAAAAACAGCGGCAGTCGTTGAACAGTCGCTGAGCCGACAGGCGCTGGCTGCACAGAAAGCGGGGATTTCCGTCGGGCAGTATAAAGCC
GCCATGCGTATGCTGCCTGCACAGTTCACCGACGTGGCCACGCAGCTTGCAGGCGGGCAAAGTCCGTGGCTGATCCTGCTGCAACAGGGGG
GGCAGGTGAAGGACTCCTTCGGCGGGATGATCCCCATGTTCAGGGGGCTTGCCGGTGCGATCACCCTGCCGATGGTGGGGGCCACCTCGCT
GGCGGTGGCGACCGGTGCGCTGGCGTATGCCTGGTATCAGGGCAACTCAACCCTGTCCGATTTCAACAAAACGCTGGTCCTTTCCGGCAATC
AGGCGGGACTGACGGCAGATCGTATGCTGGTCCTGTCCAGAGCCGGGCAGGCGGCAGGGCTGACGTTTAACCAGACCAGCGAGTCACTCAG
CGCACTGGTTAAGGCGGGGGTAAGCGGTGAGGCTCAGATTGCGTCCATCAGCCAGAGTGTGGCGCGTTTCTCCTCTGCATCCGGCGTGGAG
GTGGACAAGGTCGCTGAAGCCTTCGGGAAGCTGACCACAGACCCGACGTCGGGGCTGACGGCGATGGCTCGCCAGTTCCATAACGTGTCGG
CGGAGCAGATTGCGTATGTTGCTCAGTTGCAGCGTTCCGGCGATGAAGCCGGGGCATTGCAGGCGGCGAACGAGGCCGCAACGAAAGGGT
TTGATGACCAGACCCGCCGCCTGAAAGAGAACATGGGCACGCTGGAGACCTGGGCAGACAGGACTGCGCGGGCATTCAAATCCATGTGGGA
TGCGGTGCTGGATATTGGTCGTCCTGATACCGCGCAGGAGATGCTGATTAAGGCAGAGGCTGCGTATAAGAAAGCAGACGACATCTGGAAT
CTGCGCAAGGATGATTATTTTGTTAACGATGAAGCGCGGGCGCGTTACTGGGATGATCGTGAAAAGGCCCGTCTTGCGCTTGAAGCCGCCCG
AAAGAAGGCTGAGCAGCAGACTCAACAGGACAAAAATGCGCAGCAGCAGAGCGATACCGAAGCGTCACGGCTGAAATATACCGAAGAGGC
GCAGAAGGCTTACGAACGGCTGCAGACGCCGCTGGAGAAATATACCGCCCGTCAGGAAGAACTGAACAAGGCACTGAAAGACGGGAAAAT
CCTGCAGGCGGATTACAACACGCTGATGGCGGCGGCGAAAAAGGATTATGAAGCGACGCTGAAAAAGCCGAAACAGTCCAGCGTGAAGGT
GTCTGCGGGCGATCGTCAGGAAGACAGTGCTCATGCTGCCCTGCTGACGCTTCAGGCAGAACTCCGGACGCTGGAGAAGCATGCCGGAGCA
AATGAGAAAATCAGCCAGCAGCGCCGGGATTTGTGGAAGGCGGAGAGTCAGTTCGCGGTACTGGAGGAGGCGGCGCAACGTCGCCAGCTG
TCTGCACAGGAGAAATCCCTGCTGGCGCATAAAGATGAGACGCTGGAGTACAAACGCCAGCTGGCTGCACTTGGCGACAAGGTTACGTATCA
GGAGCGCCTGAACGCGCTGGCGCAGCAGGCGGATAAATTCGCACAGCAGCAACGGGCAAAACGGGCCGCCATTGATGCGAAAAGCCGGGG
GCTGACTGACCGGCAGGCAGAACGGGAAGCCACGGAACAGCGCCTGAAGGAACAGTATGGCGATAATCCGCTGGCGCTGAATAACGTCAT
GTCAGAGCAGAAAAAGACCTGGGCGGCTGAAGACCAGCTTCGCGGGAACTGGATGGCAGGCCTGAAGTCCGGCTGGAGTGAGTGGGAAG
AGAGCGCCACGGACAGTATGTCGCAGGTAAAAAGTGCAGCCACGCAGACCTTTGATGGTATTGCACAGAATATGGCGGCGATGCTGACCGG
CAGTGAGCAGAACTGGCGCAGCTTCACCCGTTCCGTGCTGTCCATGATGACAGAAATTCTGCTTAAGCAGGCAATGGTGGGGATTGTCGGGA
GTATCGGCAGCGCCATTGGCGGGGCTGTTGGTGGCGGCGCATCCGCGTCAGGCGGTACAGCCATTCAGGCCGCTGCGGCGAAATTCCATTTT
GCAACCGGAGGATTTACGGGAACCGGCGGCAAATATGAGCCAGCGGGGATTGTTCACCGTGGTGAGTTTGTCTTCACGAAGGAGGCAACCA
GCCGGATTGGCGTGGGGAATCTTTACCGGCTGATGCGCGGCTATGCCACCGGCGGTTATGTCGGTACACCGGGCAGCATGGCAGACAGCCG
GTCGCAGGCGTCCGGGACGTTTGAGCAGAATAACCATGTGGTGATTAACAACGACGGCACGAACGGGCAGATAGGTCCGGCTGCTCTGAAG
GCGGTGTATGACATGGCCCGCAAGGGTGCCCGTGATGAAATTCAGACACAGATGCGTGATGGTGGCCTGTTCTCCGGAGGTGGACGATGAA
GACCTTCCGCTGGAAAGTGAAACCCGGTATGGATGTGGCTTCGGTCCCTTCTGTAAGAAAGGTGCGCTTTGGTGATGGCTATTCTCAGCGAG
CGCCTGCCGGGCTGAATGCCAACCTGAAAACGTACAGCGTGACGCTTTCTGTCCCCCGTGAGGAGGCCACGGTACTGGAGTCGTTTCTGGAA
GAGCACGGGGGCTGGAAATCCTTTCTGTGGACGCCGCCTTATGAGTGGCGGCAGATAAAGGTGACCTGCGCAAAATGGTCGTCGCGGGTCA
GTATGCTGCGTGTTGAGTTCAGCGCAGAGTTTGAACAGGTGGTGAACTGATGCAGGATATCCGGCAGGAAACACTGAATGAATGCACCCGT
GCGGAGCAGTCGGCCAGCGTGGTGCTCTGGGAAATCGACCTGACAGAGGTCGGTGGAGAACGTTATTTTTTCTGTAATGAGCAGAACGAAA
AAGGTGAGCCGGTCACCTGGCAGGGGCGACAGTATCAGCCGTATCCCATTCAGGGGAGCGGTTTTGAACTGAATGGCAAAGGCACCAGTAC
GCGCCCCACGCTGACGGTTTCTAACCTGTACGGTATGGTCACCGGGATGGCGGAAGATATGCAGAGTCTGGTCGGCGGAACGGTGGTCCGG
CGTAAGGTTTACGCCCGTTTTCTGGATGCGGTGAACTTCGTCAACGGAAACAGTTACGCCGATCCGGAGCAGGAGGTGATCAGCCGCTGGCG
CATTGAGCAGTGCAGCGAACTGAGCGCGGTGAGTGCCTCCTTTGTACTGTCCACGCCGACGGAAACGGATGGCGCTGTTTTTCCGGGACGTA
TCATGCTGGCCAACACCTGCACCTGGACCTATCGCGGTGACGAGTGCGGTTATAGCGGTCCGGCTGTCGCGGATGAATATGACCAGCCAACG
TCCGATATCACGAAGGATAAATGCAGCAAATGCCTGAGCGGTTGTAAGTTCCGCAATAACGTCGGCAACTTTGGCGGCTTCCTTTCCATTAAC
AAACTTTCGCAGTAAATCCCATGACACAGACAGAATCAGCGATTCTGGCGCACGCCCGGCGATGTGCGCCAGCGGAGTCGTGCGGCTTCGTG
GTAAGCACGCCGGAGGGGGAAAGATATTTCCCCTGCGTGAATATCTCCGGTGAGCCGGAGGCGTATTTCCGTATGTCGCCGGAAGACTGGC
TGCAGGCAGAAATGCAGGGTGAGATTGTGGCGCTGGTCCACAGCCACCCCGGTGGTCTGCCCTGGCTGAGTGAGGCCGACCGGCGGCTGCA
GGTGCAGAGTGATTTGCCGTGGTGGCTGGTCTGCCGGGGGACGATTCATAAGTTCCGCTGTGTGCCGCATCTCACCGGGCGGCGCTTTGAGC
ACGGTGTGACGGACTGTTACACACTGTTCCGGGATGCTTATCATCTGGCGGGGATTGAGATGCCGGACTTTCATCGTGAGGATGACTGGTGG
CGTAACGGCCAGAATCTCTATCTGGATAATCTGGAGGCGACGGGGCTGTATCAGGTGCCGTTGTCAGCGGCACAGCCGGGCGATGTGCTGC
TGTGCTGTTTTGGTTCATCAGTGCCGAATCACGCCGCAATTTACTGCGGCGACGGCGAGCTGCTGCACCATATTCCTGAACAACTGAGCAAAC
GAGAGAGGTACACCGACAAATGGCAGCGACGCACACACTCCCTCTGGCGTCACCGGGCATGGCGCGCATCTGCCTTTACGGGGATTTACAAC
GATTTGGTCGCCGCATCGACCTTCGTGTGAAAACGGGGGCTGAAGCCATCCGGGCACTGGCCACACAGCTCCCGGCGTTTCGTCAGAAACTG
AGCGACGGCTGGTATCAGGTACGGATTGCCGGGCGGGACGTCAGCACGTCCGGGTTAACGGCGCAGTTACATGAGACTCTGCCTGATGGCG
CTGTAATTCATATTGTTCCCAGAGTCGCCGGGGCCAAGTCAGGTGGCGTATTCCAGATTGTCCTGGGGGCTGCCGCCATTGCCGGATCATTCT
TTACCGCCGGAGCCACCCTTGCAGCATGGGGGGCAGCCATTGGGGCCGGTGGTATGACCGGCATCCTGTTTTCTCTCGGTGCCAGTATGGTG
CTCGGTGGTGTGGCGCAGATGCTGGCACCGAAAGCCAGAACTCCCCGTATACAGACAACGGATAACGGTAAGCAGAACACCTATTTCTCCTC
ACTGGATAACATGGTTGCCCAGGGCAATGTTCTGCCTGTTCTGTACGGGGAAATGCGCGTGGGGTCACGCGTGGTTTCTCAGGAGATCAGCA
CGGCAGACGAAGGGGACGGTGGTCAGGTTGTGGTGATTGGTCGCTGATGCAAAATGTTTTATGTGAAACCGCCTGCGGGCGGTTTTGTCATT
TATGGAGCGTGAGGAATGGGTAAAGGAAGCAGTAAGGGGCATACCCCGCGCGAAGCGAAGGACAACCTGAAGTCCACGCAGTTGCTGAGT
GTGATCGATGCCATCAGCGAAGGGCCGATTGAAGGTCCGGTGGATGGCTTAAAAAGCGTGCTGCTGAACAGTACGCCGGTGCTGGACACTG

AGGGGAATACCAACATATCCGGTGTCACGGTGGTGTTCCGGGCTGGTGAGCAGGAGCAGACTCCGCCGGAGGGATTTGAATCCTCCGGCTC
CGAGACGGTGCTGGGTACGGAAGTGAAATATGACACGCCGATCACCCGCACCATTACGTCTGCAAACATCGACCGTCTGCGCTTTACCTTCG
GTGTACAGGCACTGGTGGAAACCACCTCAAAGGGTGACAGGAATCCGTCGGAAGTCCGCCTGCTGGTTCAGATACAACGTAACGGTGGCTG
GGTGACGGAAAAAGACATCACCATTAAGGGCAAAACCACCTCGCAGTATCTGGCCTCGGTGGTGATGGGTAACCTGCCGCCGCGCCCGTTTA
ATATCCGGATGCGCAGGATGACGCCGGACAGCACCACAGACCAGCTGCAGAACAAAACGCTCTGGTCGTCATACACTGAAATCATCGATGTG
AAACAGTGCTACCCGAACACGGCACTGGTCGGCGTGCAGGTGGACTCGGAGCAGTTCGGCAGCCAGCAGGTGAGCCGTAATTATCATCTGC
GCGGGCGTATTCTGCAGGTGCCGTCGAACTATAACCCGCAGACGCGGCAATACAGCGGTATCTGGGACGGAACGTTTAAACCGGCATACAG
CAACAACATGGCCTGGTGTCTGTGGGATATGCTGACCCATCCGCGCTACGGCATGGGGAAACGTCTTGGTGCGGCGGATGTGGATAAATGG
GCGCTGTATGTCATCGGCCAGTACTGCGACCAGTCAGTGCCGGACGGCTTTGGCGGCACGGAGCCGCGCATCACCTGTAATGCGTACCTGAC
CACACAGCGTAAGGCGTGGGATGTGCTCAGCGATTTCTGCTCGGCGATGCGCTGTATGCCGGTATGGAACGGGCAGACGCTGACGTTCGTG
CAGGACCGACCGTCGGATAAGACGTGGACCTATAACCGCAGTAATGTGGTGATGCCGGATGATGGCGCGCCGTTCCGCTACAGCTTCAGCG
CCCTGAAGGACCGCCATAATGCCGTTGAGGTGAACTGGATTGACCCGAACAACGGCTGGGAGACGGCGACAGAGCTTGTTGAAGATACGCA
GGCCATTGCCTCAGCAGCAACAACAAACAAGCAAGAAAGAACAAACAAGGAAACAAGACAAAGAACCAAAGAACACGACCTCAGCTGTTAC
GAAGATGGATGCCTTTGGCTGTACCAGCCGGGGGCAGGCACACCGCGCCGGGCTGTGGCTGATTAAAACAGAACTGCTGGAAACGCAGACC
GTGGATTTCAGCGTCGGCGCAGAAGGGCTTCGCCATGTACCGGGCGATGTTATTGAAATCTGCGATGATGACTATGCCGGTATCAGCACCGG
TGGTCGTGTGCTGGCGGTGAACAGCCAGACCCGGACGCTGACGCTCGACCGTGAAATCACGCTGCCATCCTCCGGTACCGAGCTCGAATTCA
CTGGCCGTCGTTTTACAACGTCGTGACTGGGAAAACCCTGGCGTTACCCAACTTAATCGCCTTGCAGCACATCCCCCTTTCGCCAGCTGGCGTA
ATAGCGAAGAGGCCCGCACCGATCGCCCTTCCCAACAGTTGCGCAGCCTGAATGGCGAATGGCGCCTGATGCGGTATTTTCTCCTTACGCATC
TGTGCGGTATTTCACACCGCATATGGTGCACTCTCAGTACAATCTGCTCTGATGCCGCATAGTTAAGCCAGCCCCGACACCCGCCAACACCCG
CTGACGCGCCCTGACGGGCTTGTCTGCTCCCGGCATCCGCTTACAGACAAGCTGTGACCGTCTCCGGGAGCTGCATGTGTCAGAGGTTTTCAC
CGTCATCACCGAAACGCGCGAGACGAAAGGGCCTCGTGATACGCCTATTTTTATAGGTTAATGTCATGATAATAATGGTTTCTTAGACGTCAG
GTGGCACTTTTCGGGGAAATGTGCGCGGAACCCCTATTTGTTTATTTTTCTAAATACATTCAAATATGTATCCGCTCATGAGACAATAACCCTG
ATAAATGCTTCAATAATATTGAAAAAGGAAGAGTATGAGTATTCAACATTTCCGTGTCGCCCTTATTCCCTTTTTTGCGGCATTTTGCCTTCCTG
TTTTTGCTCACCCAGAAACGCTGGTGAAAGTAAAAGATGCTGAAGATCAGTTGGGTGCACGAGTGGGTTACATCGAACTGGATCTCAACAGC
GGTAAGATCCTTGAGAGTTTTCGCCCCGAAGAACGTTTTCCAATGATGAGCACTTTTAAAGTTCTGCTATGTGGCGCGGTATTATCCCGTATTG
ACGCCGGGCAAGAGCAACTCGGTCGCCGCATACACTATTCTCAGAATGACTTGGTTGAGTACTCACCAGTCACAGAAAAGCATCTTACGGAT
GGCATGACAGTAAGAGAATTATGCAGTGCTGCCATAACCATGAGTGATAACACTGCGGCCAACTTACTTCTGACAACGATCGGAGGACCGAA
GGAGCTAACCGCTTTTTTGCACAACATGGGGGATCATGTAACTCGCCTTGATCGTTGGGAACCGGAGCTGAATGAAGCCATACCAAACGACG
AGCGTGACACCACGATGCCTGTAGCAATGGCAACAACGTTGCGCAAACTATTAACTGGCGAACTACTTACTCTAGCTTCCCGGCAACAATTAA
TAGACTGGATGGAGGCGGATAAAGTTGCAGGACCACTTCTGCGCTCGGCCCTTCCGGCTGGCTGGTTTATTGCTGATAAATCTGGAGCCGGT
GAGCGTGGGTCTCGCGGTATCATTGCAGCACTGGGGCCAGATGGTAAGCCCTCCCGTATCGTAGTTATCTACACGACGGGGAGTCAGGCAAC
TATGGATGAACGAAATAGACAGATCGCTGAGATAGGTGCCTCACTGATTAAGCATTGGTAACTGTCAGACCAAGTTTACTCATATATACTTTA
GATTGATTTAAAACTTCATTTTTAATTTAAAAGGATCTAGGTGAAGATCCTTTTTGATAATCTCATGACCAAAATCCCTTAACGTGAGTTTTCGT
TCCACTGAGCGTCAGACCCCGTAGAAAAGATCAAAGGATCTTCTTGAGATCCTTTTTTTCTGCGCGTAATCTGCTGCTTGCAAACAAAAAAACC
ACCGCTACCAGCGGTGGTTTGTTTGCCGGATCAAGAGCTACCAACTCTTTTTCCGAAGGTAACTGGCTTCAGCAGAGCGCAGATACCAAATAC
TGTTCTTCTAGTGTAGCCGTAGTTAGGCCACCACTTCAAGAACTCTGTAGCACCGCCTACATACCTCGCTCTGCTAATCCTGTTACCAGTGGCT
GCTGCCAGTGGCGATAAGTCGTGTCTTACCGGGTTGGACTCAAGACGATAGTTACCGGATAAGGCGCAGCGGTCGGGCTGAACGGGGGGTT
CGTGCACACAGCCCAGCTTGGAGCGAACGACCTACACCGAACTGAGATACCTACAGCGTGAGCTATGAGAAAGCGCCACGCTTCCCGAAGG
GAGAAAGGCGGACAGGTATCCGGTAAGCGGCAGGGTCGGAACAGGAGAGCGCACGAGGGAGCTTCCAGGGGGAAACGCCTGGTATCTTT
ATAGTCCTGTCGGGTTTCGCCACCTCTGACTTGAGCGTCGATTTTTGTGATGCTCGTCAGGGGGGCGGAGCCTATGGAAAAACGCCAGCAAC
GCGGCCTTTTTACGGTTCCTGGCCTTTTGCTGGCCTTTTGCTCACATGTTCTTTCCTGCGTTATCCCCTGATTCTGTGGATAACCGTATTACCGCC
TTTGAGTGAGCTGATACCGCTCGCCGCAGCCGAACGACCGAGCGCAGCGAGTCAGTGAGCGAGGAAGCGGAAGAGCGCCCAATACGCAAA
CCGCCTCTCCCCGCGCGTTGGCCGATTCATTAATGCAGCTGGCACGACAGGTTTCCCGACTGGAAAGCGGGCAGTGAGCGCAACGCAATTAA
TGTGAGTTAGCTCACTCATTAGGCACCCCAGGCTTTACACTTTATGCTTCCGGCTCGTATGTTGTGTGGAATTGTGAGCGGATAACAATTTCAC
ACAGGAAACAGCTATGACCATGATTACGCCAAGCTTGCATGCCTGCAGGTCGACTCTAGAGGATCCTCAACTGTGAGGAGGCTCACGGACGC
GAAGAACAGGCACGCGTGCTGGCAGAAACCCCCGGTATGACCGTGAAAACGGCCCGCCGCATTCTGGCCGCAGCACCACAGAGTGCACAGG
CGCGCAGTGACACTGCGCTGGATCGTCTGATGCAGGGGGCACCGGCACCGCTGGCTGCAGGTAACCCGGCATCTGATGCCGTTAACGATTTG
CTGAACACACCAGTGTAAGGGATGTTTATGACGAGCAAAGAAACCTTTACCCATTACCAGCCGCAGGGCAACAGTGACCCGGCTCATACCGC
AACCGCGCCCGGCGGATTGAGTGCGAAAGCGCCTGCAATGACCCCGCTGATGCTGGACACCTCCAGCCGTAAGCTGGTTGCGTGGGATGGC
ACCACCGACGGTGCTGCCGTTGGCATTCTTGCGGTTGCTGCTGACCAGACCAGCACCACGCTGACGTTCTACAAGTCCGGCACGTTCCGTTAT
GAGGATGTGCTCTGGCCGGAGGCTGCCAGCGACGAGACGAAAAAACGGACCGCGTTTGCCGGAACGGCAATCAGCATCGTTTAACTTTACC
CTTCATCACTAAAGGCCGCCTGTGCGGCTTTTTTTACGGGATTTTTTTATGTCGATGTACACAACCGCCCAACTGCTGGCGGCAAATGAGCAGA
AATTTAAGTTTGATCCGCTGTTTCTGCGTCTCTTTTTCCGTGAGAGCTATCCCTTCACCACGGAGAAAGTCTATCTCTCACAAATTCCGGGACTG
GTAAACATGGCGCTGTACGTTTCGCCGATTGTTTCCGGTGAGGTTATCCGTTCCCGTGGCGGCTCCACCTCTGAATTTACGCCGGGATATGTC
AAGCCGAAGCATGAAGTGAATCCGCAGATGACCCTGCGTCGCCTGCCGGATGAAGATCCGCAGAATCTGGCGGACCCGGCTTACCGCCGCC
GTCGCATCATCATGCAGAACATGCGTGACGAAGAGCTGGCCATTGCTCAGGTCGAAGAGATGCAGGCAGTTTCTGCCGTGCTTAAGGGCAA
ATACACCATGACCGGTGAAGCCTTCGATCCGGTTGAGGTGGATATGGGCCGCAGTGAGGAGAATAACATCACGCAGTCCGGCGGCACGGAG
TGGAGCAAGCGTGACAAGTCCACGTATGACCCGACCGACGATATCGAAGCCTACGCGCTGAACGCCAGCGGTGTGGTGAATATCATCGTGTT

CGATCCGAAAGGCTGGGCGCTGTTCCGTTCCTTCAAAGCCGTCAAGGAGAAGCTGGATACCCGTCGTGGCTCTAATTCCGAGCTGGAGACAG
CGGTGAAAGACCTGGGCAAAGCGGTGTCCTATAAGGGGATGTATGGCGATGTGGCCATCGTCGTGTATTCCGGACAGTACGTGGAAAACGG
CGTCAAAAAGAACTTCCTGCCGGACAACACGATGGTGCTGGGGAACACTCAGGCACGCGGTCTGCGCACCTATGGCTGCATTCAGGATGCG
GACGCACAGCGCGAAGGCATTAACGCCTCTGCCCGTTACCCGAAAAACTGGGTGACCACCGGCGATCCGGCGCGTGAGTTCACCATGATTCA
GTCAGCACCGCTGATGCTGCTGGCTGACCCTGATGAGTTCGTGTCCGTACAACTGGCGTAATCATGGCCCTTCGGGGCCATTGTTTCTCTGTG
GAGGAGTCCATGACGAAAGATGAACTGATTGCCCGTCTCCGCTCGCTGGGTGAACAACTGAACCGTGATGTCAGCCTGACGGGGACGAAAG
AAGAACTGGCGCTCCGTGTGGCAGAGCTGAAAGAGGAGCTTGATGACACGGATGAAACTGCCGGTCAGGACACCCCTCTCAGCCGGGAAAA
TGTGCTGACCGGACATGAAAATGAGGTGGGATCAGCGCAGCCGGATACCGTGATTCTGGATACGTCTGAACTGGTCACGGTCGTGGCACTG
GTGAAGCTGCATACTGATGCACTTCACGCCACGCGGGATGAACCTGTGGCATTTGTGCTGCCGGGAACGGCGTTTCGTGTCTCTGCCGGTGT
GGCAGCCGAAATGACAGAGCGCGGCCTGGCCAGAATGCAATAACGGGAGGCGCTGTGGCTGATTTCGATAACCTGTTCGATGCTGCCATTG
CCCGCGCCGATGAAACGATACGCGGGTACATGGGAACGTCAGCCACCATTACATCCGGTGAGCAGTCAGGTGCGGTGATACGTGGTGTTTTT
GATGACCCTGAAAATATCAGCTATGCCGGACAGGGCGTGCGCGTTGAAGGCTCCAGCCCGTCCCTGTTTGTCCGGACTGATGAGGTGCGGCA
GCTGCGGCGTGGAGACACGCTGACCATCGGTGAGGAAAATTTCTGGGTAGATCGGGTTTCGCCGGATGATGGCGGAAGTTGTCATCTCTGG
CTTGGACGGGGCGTACCGCCTGCCGTTAACCGTCGCCGCTGAAAGGGGGATGTATGGCCATAAAAGGTCTTGAGCAGGCCGTTGAATCAGC
AAAACGGCGGTGCCTGGTGCCGCCGCAATGGCCATTAACCGCGTTGCTTCATCCGCGATATCGCAGTCGGCGTCACAGGTTGCCCGTGAGAC
AAAGGTACGCCGGAAACTGGTAAAGGAAAGGGCCAGGCTGAAAAGGGCCACGGTCAAAAATCCGCAGGCCAGAATCAAAGTTAACCGGGG
GGATTTGCCCGTAATCAAGCTGGGTAATGCGCGGGTTGTCCTTTCGCGCCGCAGGCGTCGTAAAAAGGGGCAGCGTTCATCCCTGAAAGGTG
GCGGCAGCGTGCTTGTGGTGGGTAACCGTCGTATTCCCGGCGCGTTTATTCAGCAACTGAAAAATGGCCGGTGGCATGTCATGCAGCGTGTG
GCTGGGAAAAACCGTTACCCCATTGATGTGGTGAAAATCCCGATGGCGGTGCCGCTGACCACGGCGTTTAAACAAAATATTGAGCGGATACG
GCGTGAACGTCTTCCGAAAGAGCTGGGCTATGCGCTGCAGCATCAACTGAGGATGGTAATAAAGCGATGAAACATACTGAACTCCGTGCAGC
CGTACTGGATGCACTGGAGAAGCATGACACCGGGGCGACGTTTTTTGATGGTCGCCCCGCTGTTTTTGATGAGGCGGATTTTCCGGCAGTTG
CCGTTTATCTCACCGGCGCTGAATACACGGGCGAAGAGCTGGACAGCGATACCTGGCAGGCGGAGCTGCATATCGAAGTTTTCCTGCCTGCT
CAGGTGCCGGATTCAGAGCTGGATGCGTGGATGGAGTCCCGGATTTATCCGGTGATGAGCGATATCCCGGCACTGTCAGATTTGATCACCAG
TATGGTGGCCAGCGGCTATGACTACCGGCGCGACGATGATGCGGGCTTGTGGAGTTCAGCCGATCTGACTTATGTCATTACCTATGAAATGT
GAGGACGCTATGCCTGTACCAAATCCTACAATGCCGGTGAAAGGTGCCGGGACCACCCTGTGGGTTTATAAGGGGAGCGGTGACCCTTACG
CGAATCCGCTTTCAGACGTTGACTGGTCGCGTCTGGCAAAAGTTAAAGACCTGACGCCCGGCGAACTGACCGCTGAGTCCTATGACGACAGC
TATCTCGATGATGAAGATGCAGACTGGACTGCGACCGGGCAGGGGCAGAAATCTGCCGGAGATACCAGCTTCACGCTGGCGTGGATGCCCG
GAGAGCAGGGGCAGCAGGCGCTGCTGGCGTGGTTTAATGAAGGCGATACCCGTGCCTATAAAATCCGCTTCCCGAACGGCACGGTCGATGT
GTTCCGTGGCTGGGTCAGCAGTATCGGTAAGGCGGTGACGGCGAAGGAAGTGATCACCCGCACGGTGAAAGTCACCAATGTGGGACGTCC
GTCGATGGCAGAAGATCGCAGCACGGTAACAGCGGCAACCGGCATGACCGTGACGCCTGCCAGCACCTCGGTGGTGAAAGGGCAGAGCAC
CACGCTGACCGTGGCCTTCCAGCCGGAGGGCGTAACCGACAAGAGCTTTCGTGCGGTGTCTGCGGATAAAACAAAAGCCACCGTGTCGGTC
AGTGGTATGACCATCACCGTGAACGGCGTTGCTGCAGGCAAGGTCAACATTCCGGTTGTATCCGGTAATGGTGACCAAGGCCTCAGCTCGTG
TTCTGTTTCTCTCTTGTTTCTTGCTTTGTTTCTTGTCTGTTTCTGTTTGTTGTTGCTCCTCAGCCCTTGGTCCGGAGAGTCAGCGATGTTCCTGAA
AACCGAATCATTTGAACATAACGGTGTGACCGTCACGCTTTCTGAACTGTCAGCCCTGCAGCGCATTGAGCATCTCGCCCTGATGAAACGGCA
GGCAGAACAGGCGGAGTCAGACAGCAACCGGAAGTTTACTGTGGAAGACGCCATCAGAACCGGCGCGTTTCTGGTGGCGATGTCCCTGTGG
CATAACCATCCGCAGAAGACGCAGATGCCGTCCATGAATGAAGCCGTTAAACAGATTGAGCAGGAAGTGCTTACCACCTGGCCCACGGAGG
CAATTTCTCATGCTGAAAACGTGGTGTACCGGCTGTCTGGTATGTATGAGTTTGTGGTGAATAATGCCCCTGAACAGACAGAGGACGCCGGG
CCCGCAGAGCCTGTTTCTGCGGGAAAGTGTTCGACGGTGAGCTGAGTTTTGCCCTGAAACTGGCGCGTGAGATGGGGCGACCCGACTGGCG
TGCCATGCTTGCCGGGATGTCATCCACGGAGTATGCCGACTGGCACCGCTTTTACAGTACCCATTATTTTCATGATGTTCTGCTGGATATGCAC
TTTTCCGGGCTGACGTACACCGTGCTCAGCCTGTTTTTCAGCGATCCGGATATGCATCCGCTGGATTTCAGTCTGCTGAACCGGCGCGAGGCT
GACGAAGAGCCTGAAGATGATGTGCTGATGCAGAAAGCGGCAGGGCTTGCCGGAGGTGTCCGCTTTGGCCCGGACGGGAATGAAGTTATC
CCCGCTTCCCCGGATGTGGCGGACATGACGGAGGATGACGTAATGCTGATGACAGTATCAGAAGGGATCGCAGGAGGAGTCCGGTATGGCT
GAACCGGTAGGCGATCTGGTCGTTGATTTGAGTCTGGATGCGGCCAGATTTGACGAGCAGATGGCCAGAGTCAGGCGTCATTTTTCTGGTAC
GGAAAGTGATGCG

### Sequence of plasmid pMDW111

tcgcgcgtttcggtgatgacggtgaaaacctctgacacatgcagctcccggagacggtcacagcttgtctgtaagcggatgccgggagcagacaagcccgtcagggcgcgtcagcgggt
gttggcgggtgtcggggctggcttaactatgcggcatcagagcagattgtactgagagtgcaccatatgcggtgtgaaataccgcacagatgcgtaaggagaaaataccgcatcaggcg
ccattcgccattcaggctgcgcaactgttgggaagggcgatcggtgcgggcctcttcgctattacgccagctggcgaaagggggatgtgctgcaaggcgattaagttgggtaacgccagg
gttttcccagtcacgacgttgtaaaacgacggccagtgAATTCGCCTGGGTGGCTTCATTCGTTCTTTTGTTCCTTATTTTGTTCCTTACTTAGTTGGTTATTTG
CTTGTTTGGTTATTTATTTCGTTGGTTATTTGGTTAATTCCTTCTTTGCTTTCTTCATTCCTTTCTTGCTTTATTCCTTGTTTTTTTGGTTTCTTAGTT
TCCTTTTCCCTAGAGGTAGCCAAAGTCTTTGCAACTATACTTTCAGCTCTGACAAATTTGTTCTTATTACTTCTCTTTTTTTTGATTTGTTCTTCCC
TCTTTTTCTTAGCTAATTCTTGTCTTTCGATTCTAGTTCTATCAGCATTTCTTTATAAATCTATTTTTTTTTTTTTTTTCGACACAAAATGTCTATTTCT
TGGAGTGCTTACTCTTCTTTTTGTTTTTACCTTGTTTCAACTCGTTTAATCTATCAACTTTTTCCTTGATCCTTTCCAAAGATAATTTGACATCACC
TTTTTGGCACTAGGTGCCACCGATGTGGAagcttggcgtaatcatggtcatagctgtttcctgtgtgaaattgttatccgctcacaattccacacaacatacgagccggaagc
ataaagtgtaaagcctggggtgcctaatgagtgagctaactcacattaattgcgttgcgctcactgcccgctttccagtcgggaaacctgtcgtgccagctgcattaatgaatcggccaac
gcgcggggagaggcggtttgcgtattgggcgctcttccgcttcctcgctcactgactcgctgcgctcggtcgttcggctgcggcgagcggtatcagctcactcaaaggcggtaatacggtta
tccacagaatcaggggataacgcaggaaagaacatgtgagcaaaaggccagcaaaaggccaggaaccgtaaaaaggccgcgttgctggcgtttttccataggctccgcccccctgac
gagcatcacaaaaatcgacgctcaagtcagaggtggcgaaacccgacaggactataaagataccaggcgtttccccctggaagctccctcgtgcgctctcctgttccgaccctgccgctt
accggatacctgtccgcctttctcccttcgggaagcgtggcgctttctcatagctcacgctgtaggtatctcagttcggtgtaggtcgttcgctccaagctgggctgtgtgcacgaaccccc
gttcagcccgaccgctgcgccttatccggtaactatcgtcttgagtccaacccggtaagacacgacttatcgccactggcagcagccactggtaacaggattagcagagcgaggtatgta
ggcggtgctacagagttcttgaagtggtggcctaactacggctacactagaagaacagtatttggtatctgcgctctgctgaagccagttaccttcggaaaaagagttggtagctcttgatc
cggcaaacaaaccaccgctggtagcggtggtttttttgtttgcaagcagcagattacgcgcagaaaaaaaggatctcaagaagatcctttgatcttttctacggggtctgacgctcagtgg
aacgaaaactcacgttaagggattttggtcatgagattatcaaaaaggatcttcacctagatccttttaaattaaaaatgaagttttaaatcaatctaaagtatatatgagtaaacttggtc
tgacagttaccaatgcttaatcagtgaggcacctatctcagcgatctgtctatttcgttcatccatagttgcctgactccccgtcgtgtagataactacgatacgggagggcttaccatctgg
ccccagtgctgcaatgataccgcgagacccacgctcaccggctccagatttatcagcaataaaccagccagccggaagggccgagcgcagaagtggtcctgcaactttatccgcctcca
tccagtctattaattgttgccgggaagctagagtaagtagttcgccagttaatagtttgcgcaacgttgttgccattgctacaggcatcgtggtgtcacgctcgtcgtttggtatggcttcatt
cagctccggttcccaacgatcaaggcgagttacatgatcccccatgttgtgcaaaaaagcggttagctccttcggtcctccgatcgttgtcagaagtaagttggccgcagtgttatcactca
tggttatggcagcactgcataattctcttactgtcatgccatccgtaagatgcttttctgtgactggtgagtactcaaccaagtcattctgagaatagtgtatgcggcgaccgagttgctcttg
cccggcgtcaatacgggataataccgcgccacatagcagaactttaaaagtgctcatcattggaaaacgttcttcggggcgaaaactctcaaggatcttaccgctgttgagatccagttcg
atgtaacccactcgtgcacccaactgatcttcagcatcttttactttcaccagcgtttctgggtgagcaaaaacaggaaggcaaaatgccgcaaaaaagggaataagggcgacacggaa
atgttgaatactcatactcttcctttttcaatattattgaagcatttatcagggttattgtctcatgagcggatacatatttgaatgtatttagaaaaataaacaaataggggttccgcgcaca
tttccccgaaaagtgccacctgacgtctaagaaaccattattatcatgacattaacctataaaaataggcgtatcacgaggccctttcgtc
